# Induction of an alternative 5’ leader enhances translation of *Inpp5e* and resistance to oncolytic virus infection

**DOI:** 10.1101/522607

**Authors:** Huy-Dung Hoang, Tyson E. Graber, Jian-Jun Jia, Nasana Vaidya, Victoria Gilchrist, Wencheng Li, Christos G. Gkogkas, Maritza Jaramillo, Seyed Mehdi Jafarnejad, Tommy Alain

## Abstract

Residual cell-intrinsic innate immunity in cancer cells hampers infection with oncolytic viruses. mRNA translation is an important component of innate immunity, yet the targeted cellular mRNAs remain ill-defined. We characterized the translatome of resistant murine “4T1” breast cancer cells infected with three of the most clinically advanced oncolytic viruses: Herpes Simplex virus 1, Reovirus and Vaccinia virus. Common among all three infections were translationally de-repressed mRNAs involved in ciliary homeostasis including *Inpp5e*, encoding an inositol 5-phosphatase that modifies lipid second messenger signalling. Translationally repressed in the uninfected condition, viral infection induced expression of an *Inpp5e* mRNA variant that lacks repressive upstream open reading frames (uORFs) within its 5’ leader and is consequently efficiently translated. Furthermore, we show that INPP5E contributes to antiviral immunity by altering virus attachment. These findings uncover a role for translational control through alternative 5’ leader expression and assign ciliary proteins such as INPP5E to the cellular antiviral response.

## INTRODUCTION

Mammalian cells possess a sophisticated cell-intrinsic innate antiviral program that is activated upon infection. Transcriptional induction of type I interferon expression (IFN-α and β) and downstream interferon stimulated genes (ISGs) is a major, well-characterized arm of the innate immune response to infection ^1,2^. Another essential but less characterized feature of this response is the mRNA translation arm of innate immunity – a reprogramming of protein synthesis to permit expression of cellular antiviral proteins while concurrently thwarting production of viral proteins ^3,4^.

Translation initiation can be modulated by several eukaryotic initiation factors (eIF) and RNA binding proteins (RBP) ^5,6^. In addition to the m^7^G cap structure which helps recruit eIFs, other cis-acting sequence elements that lie within 5’ leaders, such as 5’ terminal oligopyrimidine (TOP) motifs, upstream open reading frames (uORF), internal ribosome entry sites (IRES), RBP binding sites or localized secondary structure can govern the translational efficiency (TE) of mRNAs ^6,7^. During infection, signalling cascades that feed into mRNA translation such as PI3K/mTORC1/(S6K or 4E-BP) and ERK/MNK/eIF4E were shown to enhance translation of antiviral mRNAs including *IRF7* and *ISG15*^8–10^. Conversely, translation initiation can be transiently suppressed following infection, by preventing efficient ribosome assembly through activation of eIF2α kinases and subsequent phosphorylation of the alpha-subunit of eIF2 (P-eIF2α) ^3^. Paradoxically, some cellular mRNAs which have uORFs (e.g. *ATF4* ^11^) and/or IRES (e.g. cIAP1/*BIRC2* ^12^) in their 5’ leaders display enhanced TE in conditions of high phospho-eIF2α. Alternative mRNA transcription, splicing and polyadenylation can also indirectly modify translational output by altering 5’ leaders and 3’ untranslated regions (UTR), thus changing the composition of sequence elements that affect TE ^13–15^.

Viruses use a plethora of methods to maximize TE of their own mRNAs, from evolving 5’ leaders that are better substrates for translation due to the presence of IRES, to deploying proteins that shutdown global cellular translation (host-shutoff)^4^. Surveying which cellular and viral mRNAs are translated in this highly dynamic environment has been the subject of several recent studies ^16,17^. However, these investigations did not specifically address how translation of mRNAs encoding putative pro-or antiviral effectors might modulate infection. Furthermore, identification of host mRNAs under translation control during infection could provide targetable strategies to improve antiviral therapies or alleviate viral resistance, an undesirable feature of tumour cells in the context of oncolytic virus therapy, which represents a promising class of cancer therapeutics that relies on natural or engineered cancer cell tropism and mobilization of adaptive, anti-tumour immunity ^18,19^.

In this work, we queried which specific substrates of translation contribute to the viral resistance of 4T1 breast cancer cells with each of three leading oncolytic viruses: Herpes Simplex Virus 1-1716 from Sorrento Therapeutics (HSV1), Reovirus Type 3 Dearing - “Reolysin” from Oncoytics Biotech (Reovirus), and Vaccinia Virus “JX-594” from Sillajen (formerly Jennerex Biotherapeutics) (VACV). Comparing viral versus mock infected, we identified translationally upregulated host mRNAs common to all three infections. We show that these mRNAs, whose 5’ leader are enriched in uORFs, are translationally repressed in mock condition and become de-repressed upon infection. Interestingly, this subset includes mRNAs encoding proteins associated with primary cilium homeostasis. We characterized the important ciliopathy gene *Inpp5e*, encoding an inositol 5- phosphatase, and describe a virus-induced RNA variant switch that releases its uORF-mediated translation repression. This response limits viral propagation as cells deficient in INPP5E exhibit increased cell surface attachment of virions and subsequent infection efficiency. Together, these findings highlight the dynamic landscape of alternative 5’ leader usage during viral infection and identify INPP5E as a translationally induced antiviral effector that limits oncolytic virus efficacy.

## RESULTS

### Distinct transcriptional and translational host responses to oncolytic viruses

To assess for translationally regulated innate immune genes, we used HSV1-1716 (HSV1), Reolysin (Reovirus), and JX-594 (VACV) individually to infect 4T1 cells, a murine mammary carcinoma model that is refractory to viral oncolysis and closely resembles stage IV human breast cancer ^20,21^. Each of these viruses has a different rate of infection that eventually results in the shutdown of host cell translation ^22,23^. We therefore selected an effective dose that is cytopathic for 50% (ED_50_) of 4T1 cells at 48 hours post-infection (**Fig. S1A**). At 18 hours post-infection, polysome profiles (**Fig. 1A**) and ^35^S-methionine labelling (**Fig. 1B**) showed robust viral protein synthesis while that of the host cells was only slightly affected. The majority of cells were infected at this dose and timepoint, confirmed by co-expression of a virion-derived GFP transgene in the case of HSV1 and VACV (**Fig. S1B**).

**Figure 1.**
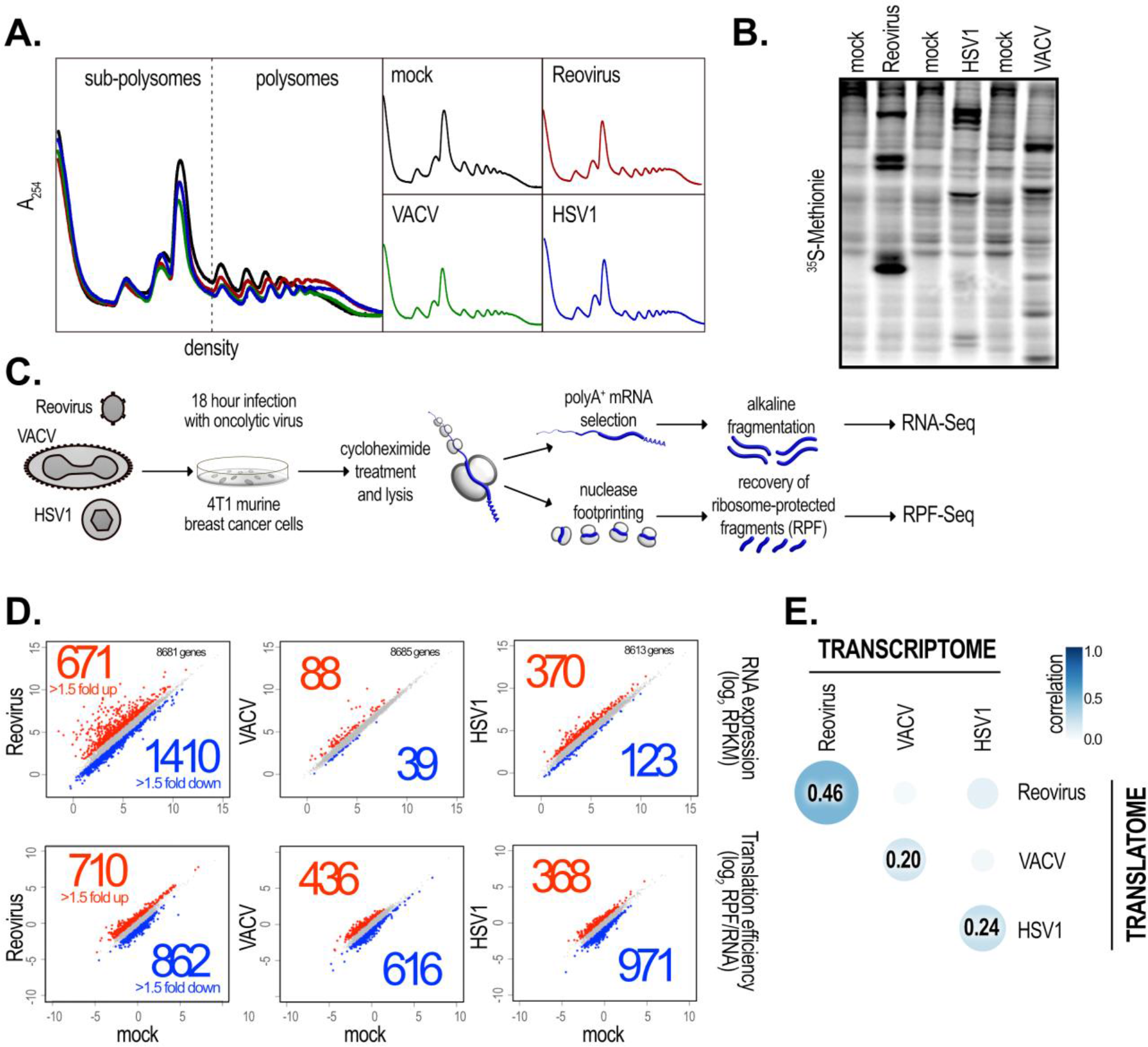
Generating the transcriptome and translatome profiles of breast cancer cells undergoing viral infection. (**A**) Polysome profiling of virus-vs. mock-infected 4T1 cells at 18 hours post-infection, or (**B**) metabolic labelling with ^35^S-methionine followed by SDS-PAGE to resolve labelled, nascent peptides, showed efficient viral translation and no global changes in cellular translation. (**C**) Schematic illustration of the ribosome profiling strategy used in this study. (**D**) Differential expression of sequenced genes (total number shown in top right of plots) at both the transcriptional and translational genome levels. The average RNA expression or translation efficiency from two biological replicates is shown. Genes that were up-or downregulated more than 1.5-fold were considered to be differentially expressed. (**E**) Correlation analysis of RNA (transcriptome) and RPF (translatome) normalized abundance between viruses (all relative to mock-infected). Pearson’s correlation coefficients are presented.

We profiled both the transcriptome and translatome signatures of 4T1 cells with each individual infection compared to mock-infected controls using the ribosome profiling method (summarized in **Fig. 1C**) ^24^. By in-parallel sequencing of ribosome-protected footprints (RPF) and total mRNA (RNA), ribosome occupancy and thus translation efficiency (TE; RPF/RNA) of individual mRNA species was quantitated ^25^. In contrast to total RNA read densities, which were constant throughout exonic regions, RPF densities were found to increase within annotated coding sequences (CDS) relative to 5’ leaders and 3’ UTRs, consistent with ribosomes engaged in translation (**Fig. S1C**). Regression analysis of reads normalized to CDS length and sequencing depth (reads per kilobase of CDS per million reads sequenced; RPKM) in two biological replicates showed a high degree of correlation at both the RNA (i.e., transcriptome) and RPF (i.e., translatome) genomic levels (**Fig. S1D**). Together, these data demonstrate that ribosome profiling successfully captured the transcriptional and translational states of the 4T1 genome following challenge with three distinct oncolytic viruses.

We next determined the differentially expressed genes (DEGs) at the transcriptome and translatome levels in virus-vs. mock-infected 4T1 cells (**Table S1**). We used a cut-off of 1.5-fold change in expression up or down and found that Reovirus modified transcription to a higher degree (24% of sequenced genes) than HSV1 (5.7%) or VACV (1.5%) (**Fig. 1D, top**). In stark contrast, TEs were perturbed more consistently between all three viruses: Reovirus (18% of sequenced genes), VACV (12%) and HSV1 (15%) (**Fig. 1D, bottom**). We also found a poor correlation between the transcriptome and the translatome (Pearson’s correlation coefficients of 0.46, 0.20, and 0.24 for Reovirus, VACV and HSV1 vs. mock, respectively) (**Fig. 1E**), which confirmed as previously reported in other studies that changes in the transcriptional profile of any given gene is a poor predictor of its TE in this context ^26–29^.

### Genes in the shared infected-translatome function in pathways not previously associated with viral infections

A primary objective of this study was to identify innate immunity effectors that could function during all three infections as a general antiviral program. To this end, we determined the common set of DEGs at both the transcriptome and translatome levels (**Fig. 2A and Table S2**). We performed Gene Ontology (GO) functional enrichment analysis on these shared sets and found the expected functional groups at the transcriptional level, including terms encompassing pattern recognition signalling, response to infection, inflammatory response, and nucleic acid binding (**Fig. 2B**). Of note, many previously validated ISGs ^1^ were found to be uniquely upregulated at the level of transcription with Reovirus, VACV and HSV1 infections, respectively (**Fig. S2A**). Moreover, these ISGs populated 38% (13/34) of the transcriptionally upregulated DEGs common to all 3 infections (**Fig. S2B**).

**Figure 2.**
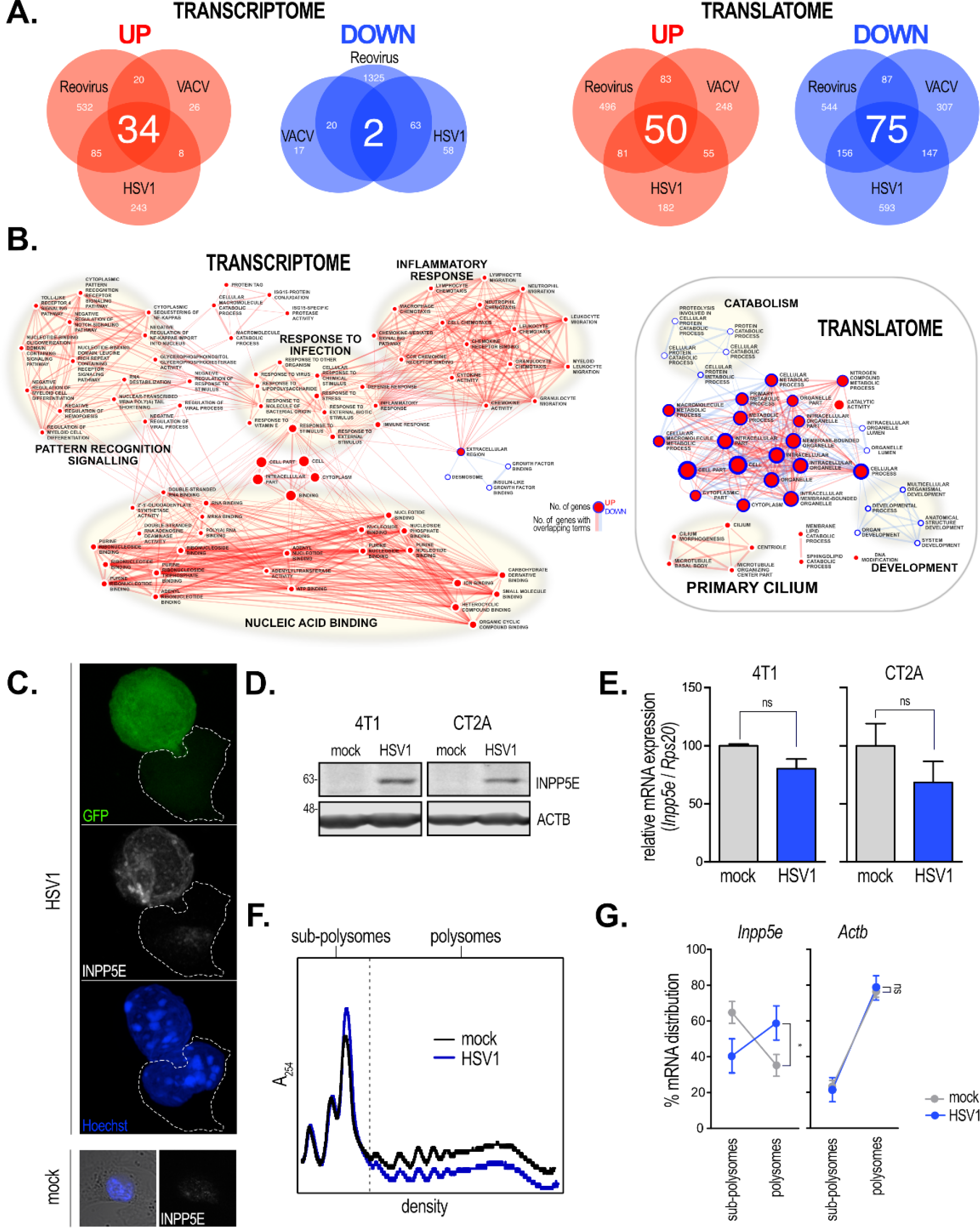
The transcriptional and translational responses to oncolytic viruses are distinct. (**A**) Venn diagrams illustrating the common/shared and unique sets of DEGs between viral infections at the transcriptional and translational levels. (**B**) Network analysis of enriched GO terms for the DEGs commonly regulated by all three oncolytic viruses at both genomic levels. Generic terms are concentrated at the centre and specific terms lie at the extremities. (**C**) Representative confocal images of anti-INPP5E-stained 4T1 cells infected with HSV1-GFP or not (mock, lower panel). An uninfected cell is outlined by a dashed line and displays poor INPP5E staining. Hoechst staining indicates the location of the nucleus. (**D**) Representative western blot of steady-state INPP5E protein expression in mock- and HSV1- infected 4T1 or CT2A cells at the 18 hours post-infection at 1 MOI. ACTB is included as a loading control. Apparent molecular weights are indicated to the left in kDa (**E**) qRT-PCR of Inpp5e mRNA abundance normalized to that of Rps20 in mock- or HSV1-infected cells at 24 hours post-infection at MOI of 5 in 4T1 and CT2A cells. (**F**) Representative polysome profiles of mock- and HSV1-infected CT2A cells 4 hours post-infection at MOI of 5. (**G**) Distribution of Actb (Left) and Inpp5e (Right) mRNAs across sub-polysomes (low TE) and polysomes (high TE) in mock- vs. HSV1-infected samples. Data are mean ± sem from five independent experiments.

Critically, functional enrichment at the level of the shared infected-translatome was found to be very different, with few specific GO terms. Translationally downregulated genes fell into categories encompassing catabolic and developmental cellular functions, while the upregulated common set was found to be enriched in genes involved in microtubule organization and the primary cilium, an organelle with a unique cytoskeleton and sub-cellular proteome that is often referred to as the cell’s signaling antenna ^30^ (**Fig. 2B, Table S3**). Confirming the GO analysis, there was significant enrichment (p=1.25×10^−4^) of genes encoding proteins that constitute the ciliary interactome (**Fig. S2C**) ^31^. Thus, using gene expression profiling, anticipated transcriptional but unexpected translational signatures of the antiviral state in oncolytic virus-infected 4T1 were obtained. This suggests that viral infection engenders a distinctive reprogramming of cellular translation.

### Inpp5e is translationally de-repressed upon oncolytic virus infection

We next aimed to validate the increased TE of the primary cilium genes identified in the network analysis. Two of these include the important ciliopathy genes *Inpp5e*, an inositol 5-phosphatase linked to Joubert Syndrome, and *Bbs9*, encoding a protein of unknown activity associated with Bardet-Biedel Syndrome (BBS) that regulates transport of ciliary proteins ^32^. We observed that INPP5E protein appeared at low level in mock condition but is dramatically induced particularly with HSV1 infection in 4T1 cells 18 hours post-infection (**Fig. 2C**). This induced protein is localized to both the cell periphery and possibly centriole regions, a localization in agreement with previous studies ^33^ (**Fig. 2C**). Consistent with immunofluorescence results, western blotting demonstrated that INPP5E is barely detectable in 4T1 cells in control conditions, but notably increased upon HSV1 infection, (**Fig. 2D, left panel**). This effect was not unique to this cancer cell line as a similar response was observed in CT2A mouse glioblastoma cells (**Fig. 2D, right panel**). This induction of INPP5E protein was a post-transcriptional effect, as levels of *Inpp5e* mRNA actually trended downwards in both cell lines at 18 hours post-infection compared to control condition (**Fig. 2E**). Importantly, the other ciliary protein, BBS9, was seen to be similarly induced as detected by immunofluorescence (**Fig. S2D**).

To address the possibility that the increased protein expression of INPP5E was due to enhanced TE and not to an increase in protein stability, we determined the TE of *Inpp5e* mRNA using conventional polysome profiling, where mRNAs are resolved based on the number of associated ribosomes (**Fig. 2F**). Unlike *Actb* mRNA, of which the majority (~80%) was found in the polysome fraction in uninfected CT2A cells, *Inpp5e* resided mainly (~60%) in sub-polysomes, indicative of a repressed translation state (**Fig. 2G, left panel**). However, four hours post-HSV1 infection at high multiplicity of infection (MOI of 5), the distribution of *Inpp5e* in active polysomes positively shifted from 35.24 ± 6.12% to 59.21 ± 9.53% while that of *Actb* did not appreciably change between mock- vs. HSV1-infected cells (76.49 ± 3.2% and 78.47 ± 6.8%, respectively). As for *Bbs9*, a similar increase of mRNA distribution to the active polysome fractions, from 64.87 ± 1.96% in mock to 75.46 ± 1.44% in HSV1-infected cells was observed (**Fig. S2E**). Thus, these data demonstrate that *Inpp5e* and *Bbs9* mRNAs are under positive translational control during viral infection in cancer cells despite that increased 80S monosome and decreased polysome peaks are observed in the polysome profile suggestive of a general repression of global translation (**Fig. 2F**)

### Oncolytic viruses de-repress translation of host mRNAs enriched in uORFs

Translational control is mediated by a constellation of trans-acting RBPs and microRNAs (miRNAs) that interact with sequence and/or structural elements on mRNA substrates ^34^. To determine if translationally regulated transcripts are enriched in any particular cis-acting element(s), we queried for enrichment of trans-acting factor binding sites in annotated 5’ leaders or 3’ UTRs using the Analysis of Motif Enrichment (AME) algorithm (part of the MEME suite ^35^) and the CISBP RNA database ^36^. Using a conservative cut-off (p<0.01), the 5’ leaders were found to be enriched in SRSF9 and SRSF1 consensus binding sequences (**Fig. S3A, top**), two known mediators of RNA splicing and translation ^37^. Interestingly, different viruses are known to alter the splicing machinery of the infected cells ^38,39^. 3’ UTR enriched motifs were numerous and included KHDRB, ELAVL1 (HuR), RBMS3, HNRNPL, HNRNPLL, ENOX1 and IGF2BP3 (**Fig. S3A, bottom**). We also analyzed the network of potential miRNA binding sites using miRNet which incorporates miRNA-target mRNA interaction data from 11 databases ^40^. This revealed a number of potential miRNAs that may regulate several of the common transcripts (**Fig. S3B**). However, no single miRNA was found to target more than three transcripts, suggesting that miRNA regulation might not readily explain the selective translational control seen during infection. TE can be modified by more general sequence features such as GC content, which predicts the potential for RNA secondary structure in 5’ leaders and thus repression of translation initiation ^41^ and is negatively correlated with transcript length ^42,43^. We surveyed the GC content of these annotated transcripts as well as the lengths of their 5’ leaders, CDS and 3’ UTRs, comparing them to all mouse mRNAs that populate the NCBI Reference Sequence (RefSeq) database. While we found no significant difference in GC content, the shared translationally regulated mRNAs (either up- or downregulated) tended to have shorter 5’ leaders, CDS and 3’ UTRs (**Fig. 3A**).

**Figure 3.**
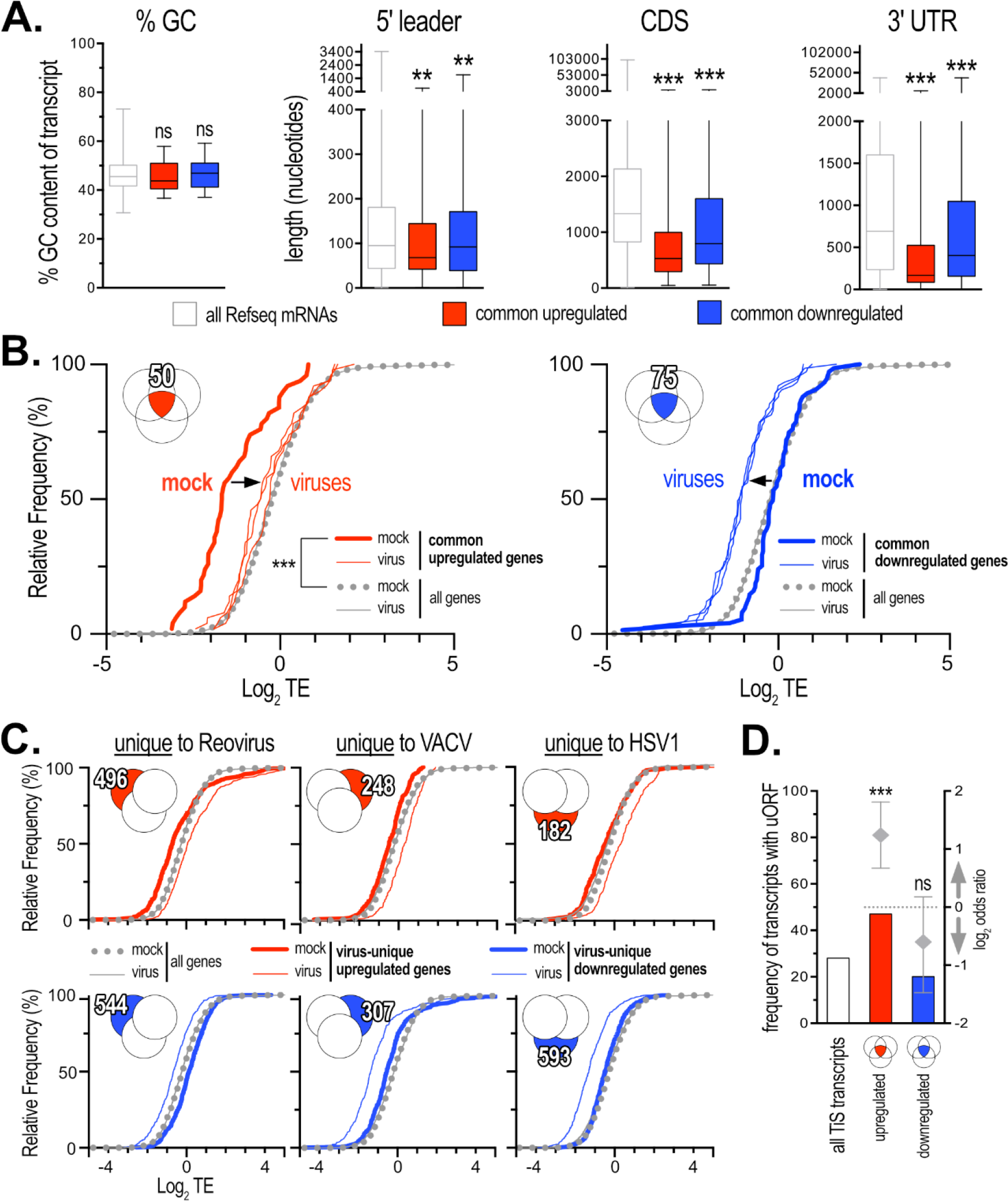
Commonly upregulated genes are de-repressed at the level of translation with infection. (**A**) Analysis of RefSeq mRNA sequence characteristics that populate the common translationally regulated gene sets compared to the entire RefSeq database (see Materials and Methods for details). Statistical significance was determined using Kruskal-Wallis with Dunn’s post-hoc testing. (**B**) Relative frequency distributions of TE for common upregulated (left panel) vs. downregulated genes (right panel) show that upregulated genes are translationally de-repressed (directional shifts indicated by an arrow). A Mann-Whitney statistical test was used. (**C**) Analysis as in **B**, but for the translationally regulated gene sets unique to each virus. (**D**) Frequency of uORFs in all TiS database mouse and human transcripts (freq.=7361/26735; http://tisdb.human.cornell.edu) compared to that present in the commonly upregulated (52/110) and downregulated (17/85) sets of transcripts. The increased frequency of uORFs in the commonly upregulated set was found to be significant by Fisher’s exact test and a log odds ratio is presented indicating overrepresentation of uORF-containing transcripts in the upregulated gene set.

In analyzing the distribution of TEs for the commonly upregulated set of genes, we noted that this subset was dramatically repressed in mock-infected cells compared to the entire sequenced set (**Fig. 3B**; compare “mock” distributions). This property appeared to be mainly due to lower RPF expression in the upregulated genes (**Figure S3C,**leftmost panel). Strikingly, viral infection acted to significantly de-repress the translation of these genes independently of changes at the RNA level (**Fig. 3B;**shift in distribution denoted by arrow, and **Fig. S3C**). In contrast, the downregulated set of genes followed a mock-infected TE distribution profile indistinguishable from the entire set of sequenced genes that shifted to lower TE upon viral infection (**Fig. 3B**, right; denoted by arrow). Furthermore, a similar analysis performed on the upregulated gene sets uniquely associated with each of the three viruses showed no clear de-repression profile suggesting that this property is a specific and common cellular response to different viral infections (**Fig. 3C**, top panels). Downregulated members of virus-unique sets behaved similarly to the shared set with respect to their TE distributions (**Fig. 3C**, bottom panels).

This repression/de-repression phenomenon has previously been ascribed to cis-acting mechanisms of translational control involving uORFs. We used an empirically-derived list of uORF-containing mouse and human mRNAs ^44,45^ and found an over-representation in the commonly upregulated set, while no enrichment was seen for the downregulated set (**Fig. 3D**). Thus, three different viruses were found to commonly affect translation of transcripts whose 5’ leaders are more likely to contain uORFs, suggesting a universal cellular response to viruses that targets cellular mRNAs harbouring uORFs.

### Inpp5e is strongly repressed through multiple uORFs which are removed following infection by expression of an alternative 5’ leader

Our data demonstrated that *Inpp5e* translation is normally repressed in 4T1 and CT2A cells, as evidenced by its low TE in mock-infected cells from ribosome profiling, polysome profiling, and the low steady-state protein expression observed by western blotting/immunofluorescence. Furthermore, the same assays showed that viral infection de-represses *Inpp5e* translation. We therefore hypothesized that this repression/de-repression shift is due to uORF-dependent translational control of *Inpp5e* mRNA. The mouse *Inpp5e* mRNA has two 5’ leader variants, the “long” and the “short”, the latter resulting from a downstream alternative transcription start site (altTSS) and alternative splicing, removing an intron that lies entirely within the 5’ leader (**Fig. S4A**). A previous genome-wide study of initiating ribosomes listed a putative uORF in both the long and short *Inpp5e 5’* leaders (**Fig. 4A**, top schema shows the long and short *Inpp5e* mRNA variant with the location of the putative uORF in dark blue) ^45^. Critically, in uninfected 4T1 cells, RPF read densities were concentrated in the *Inpp5e* leader region (**Fig. 4A**, leader region highlighted in light blue). This density shifted to the main ORF (mORF) upon HSV1, Reovirus and VACV infections, strongly suggestive of a regulatory uORF.

**Figure 4.**
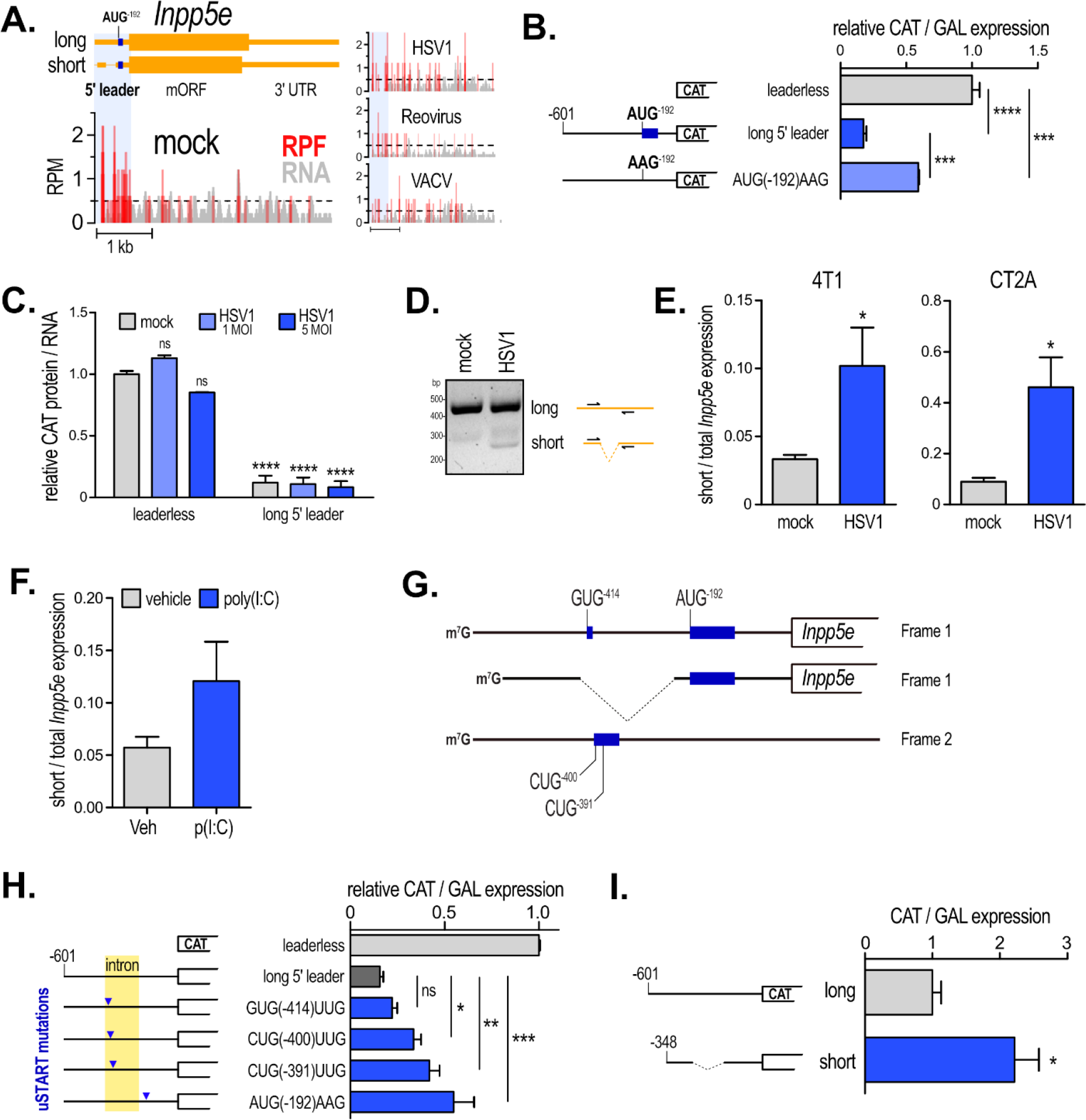
HSV1 infection induces expression of an alternative Inpp5e 5’ leader that improves TE through exclusion of repressive uORFs. (**A**) Chromatogram of read densities (reads per million, RPM) merged from two biological replicates for the Inpp5e locus at both RNA and RPF gene expression levels showing a shift of RPF reads in the 5’ leader to the mORF with infection. (**B**) Translation reporter assay in 4T1 cells showing that the long Inpp5e 5’ leader strongly represses translation of the CAT ORF. Constructs used are pictured to the left and were co-transfected with a β-galactosidase (GAL) reporter to normalize any differences in transfection. The long leader-mediated repression is substantially weakened with a single mutation that abolishes an uORF start codon (AUG[-192]AAG) demonstrating a bona fide uORF mechanism of translational regulation. (**C**) Translation reporter assay as in **B**, but in 4T1 cells transfected with in vitro transcribed CAT RNA evaluated at 18 hours post-infection. Data represent two independent experiments. (**D**) Agarose gel with resolved PCR amplicons from a 4T1 cDNA library indicate expression of both the short and long Inpp5e 5’ leaders in HSV1-infected cells (a single set of primers flank the 5’ leader intron, right). (**E**) qRT-PCR of short vs. total Inpp5e mRNA in 4T1 and CT2A cells showing increased expression of the short variant with HSV1 infection after 48 hours of infection. Data are from 5-6 independent experiments. (**F**) qRT-PCR short vs. total Inpp5e mRNA expression (right) in CT2A cells treated with poly(I:C) at the indicated times. Data are from 4 independent experiments. (**G**) Schematic of the Inpp5e 5’ leader region in both the long (top) and short (middle) mRNA transcript variants in reading frame 1 which encodes INPP5E, and frame 2 which is non-coding (bottom). Putative start codons are indicated with numbering relative to the first A of the Inpp5e AUG start codon in the long variant (NM_033134.3). Locations of potential uORFs are indicated with dark blue rectangles. (**H**) CAT reporter assays were performed as in **B** with uORF start codon mutations (GUG[-414]UUG, CUG[-400]UUG, CUG[-391]UUG, AUG[-192]AAG; indicated by blue triangles) in the long Inpp5e 5’ leader construct. (**I**) Translation reporter assay of DNA-transfected 4T1 cells showing that the short Inpp5e 5’ leader variant (348 nt) is a markedly better substrate for translation than the long 5’ leader variant (601 nt).

To determine the presence of a *bona fide* uORF, we constructed a heterologous DNA reporter plasmid consisting of the long (601 nt) *Inpp5e* leader inserted in front of a mORF encoding chloramphenicol acetyl transferase (CAT) ^12^. We found that the long *Inpp5e* leader confers a very strong repressive effect on CAT expression, with levels approaching only 20% of that observed in 4T1 cells transfected with a leaderless construct (**Fig. 4B**). Importantly, mutating the methionine-coding AUG start codon of the putative uORF at −192 nt (uORF^−192^; present in both long and short leaders) to AAG rescued CAT expression to approximately 60% of the leaderless construct (**Fig. 4B**). Together, these data show that *Inpp5e* leaders harbour an uORF whose translation represses that of the downstream cistron. This likely explains why this transcript is normally repressed at the level of translation as seen in the ribosome and polysome profiling experiments.

We next tested if *Inpp5e* uses a classical uORF de-repression mechanism to modulate its translation during viral infection. In this model, translation of uORF^−192^ would dominate over the mORF in basal conditions, while the inverse would prevail with viral infection. 4T1 cells transfected with *in vitro* synthesized, capped and polyadenylated CAT reporter RNA, however, showed no changes in *Inpp5e* leader-dependent translation with HSV1 infection even at high MOI (**Fig. 4C**). Thus, we were unable to demonstrate that the induction of *Inpp5e* translation during viral infection employs a classical uORF de-repression mechanism. An alternative possibility is that infection induces a shift in 5’ leader expression, potentially favouring the short variant of *Inpp5e*. By virtue of its lower 5’ leader complexity this transcript could conceivably display increased TE. This transcript, harbouring a shorter 3’ UTR and encoding a C-terminally truncated protein isoform, is supported in the aggregated RNA-Seq database by intron-spanning reads that account for approximately 5% of all *Inpp5e* transcripts (**Fig. S4A**), as well as a single RIKEN clone isolated from neonatal kidney tissue. These data suggest that this alternatively-transcribed and -spliced transcript is a minor player in most contexts. Notwithstanding this, we observed a decrease in the *Inpp5e* 5’ leader/CDS ratio of RNA read densities during virus infection, suggesting a shift in expression to the shorter leader variant (**Fig. S4B**). We asked whether a change in alternative splicing was a general feature of the infected 4T1 cells. Using the Mixture of Isoform (MISO) algorithm ^46^ to quantitate changes in alternative splicing from our RNA-Seq experiments, an altered splicing landscape during infection with all three viruses was found. We detected 195, 229 and 289 differentially spliced events in Reovirus-, VACV- and HSV1- vs. mock-infected 4T1 cells, respectively (**Fig. S4C, D**). However, MISO was unable to call differentially spliced events in the list of shared translationally regulated genes, potentially due to their low mapping density as most were found to be rarer mRNAs (**Fig. S3C, left**).

To monitor the expression of the short and long variants, we designed PCR primers to flank the intron in the 5’ leader and could readily detect the long variant in both mock-and HSV1-infected 4T1 cells, while the short variant was only evident in the latter condition (**Fig. 4D**). We next repeated the experiment in both 4T1 and CT2A cells using qRT-PCR primers designed to span the exon-exon junction of the short 5’ leader and found that the short variant in uninfected 4T1 or CT2A cells represented 3.32 ± 0.325% and 8.89 ± 1.53%, respectively, of the total abundance of *Inpp5e* mRNA (total *Inpp5e* was detected with primers annealing to a 3’ exon common to both the long and short transcripts) (**Fig. 4E**). Importantly, infection with HSV1 caused a significant increase in the expression of the short variant relative to total *Inpp5e* mRNA levels to 10.19 ± 2.8% and 46.08 ± 11.74% in 4T1 and CT2A, respectively (**Fig. 4E**). Moreover, poly(I:C), which mimics exposure to viral dsRNA and triggers innate immunity in part by activating PKR, was found to increase short variant expression in CT2A at 8 hours post-treatment, albeit to a lesser degree than with HSV1 (**Fig. 4F**). PKR activation has been previously shown to modulate mRNA splicing ^47^; therefore we asked if induction of the short *Inpp5e* variant was dependent on this kinase. Although we could re-capitulate the poly(I:C)-mediated induction of the short variant in wildtype mouse embryonic fibroblasts (MEFs), there was a similar induction in their PKR-null counterparts ^48,49^, suggesting that the short variant induction is PKR- independent (**Fig. S4F**).

We reasoned that there might be other uORFs in the 5’ leader intronic region that repress translation of the mORF and are spliced-out in the short 5’ leader during viral infection. Two other ribosome profiling studies in lipopolysaccharide-treated dendritic cells ^50^ and SOX-treated keratinocytes ^51^ showed relatively strong initiating ribosome peaks at two near-cognate CUG start codons, as well as a strong elongating ribosome peak at a near-cognate GUG at 400, 391 and 414 nt upstream of the main CDS, respectively (referred to here as CUG^−400^, CUG^−391^, GUG^−414^) (**Fig. S4G**). The putative uORF starting at GUG^−414^ is 48 bp and in-frame with both uORF^−192^ and the mORF; while the putative uORFs starting at CUG^−400^ and CUG^−391^ are 141 nt and 132 nt, respectively, and in-frame with each other but out-of-frame with the other uORFs and the mORF (**Fig. 4G**). To determine if these codons are used to initiate translation of *bona fide* uORFs, we mutated each of them in the *Inpp5e* long leader CAT construct and determined CAT expression as previously. Mutating the −400 and −391 CUG codons to UUG resulted in a significant enhancement of CAT expression, while mutating the −414 GUG codon to UUG slightly enhanced CAT expression, albeit not statistically significant (**Fig. 4H**). Thus, near-cognate CUG start codons in the 5’leader intron contribute to uORF-mediated repression of *Inpp5e* translation.

Finally, we asked if the short *Inpp5e* transcript variant is a better substrate for translation. Comparison of CAT expression in transfected 4T1 cells revealed that the short (spliced) 5’ leader confers a 2-fold enhancement of CAT expression relative to the long (unspliced) 5’ leader (**Fig. 4I**). Together, these data attach a complex expression profile for *Inpp5e*; repressive uORFs prevent inappropriate translation of *Inpp5e* under normal conditions but are removed following viral infection to enhance its translation.

### INPP5E acts as an antiviral effector that modifies cell attachment

We next investigated the potential function of INPP5E in modulating viral infection. We first knocked- down *Inpp5e* expression in CT2A using a short hairpin RNA (shRNA) and found an increase in HSV1 protein expression relative to cells expressing non-targeting (shNTC) RNAi as evaluated by western blot (**Fig. S5A**). We turned to CRISPR/Cas9 technology ^52^ to generate two clones of 4T1 cells depleted of INPP5E (*Inpp5e*^CRISPR#1^, *Inpp5e*^CRISPR#2^). Tracking of Indels by DEcomposition (TIDE) analysis ^53^ determined that the *Inpp5e* locus was modified at a frequency close to 100% with a mixture of 3 indels that would be predicted to disrupt expression (**Fig. S5B-D**). Western blot of 4T1 *Inpp5e*^CRISPR#1^ infected with HSV1 at a saturation MOI (of 5) shown a lack of INPP5E induction compared to wild-type 4T, confirming the knockout of the protein (**Fig. 5A**). As with the CT2A knock-down cells, *Inpp5e*^CRISPR^ cells also demonstrated increased expression of virally-expressed GFP with HSV1 (**Fig. 5B**). Interestingly, infection with VSV∆51 (a (-)ssRNA virus and another oncolytic virus candidate) was also enhanced in these cells, although no clear difference was observed with VACV (**Fig. 5B**).

**Figure 5.**
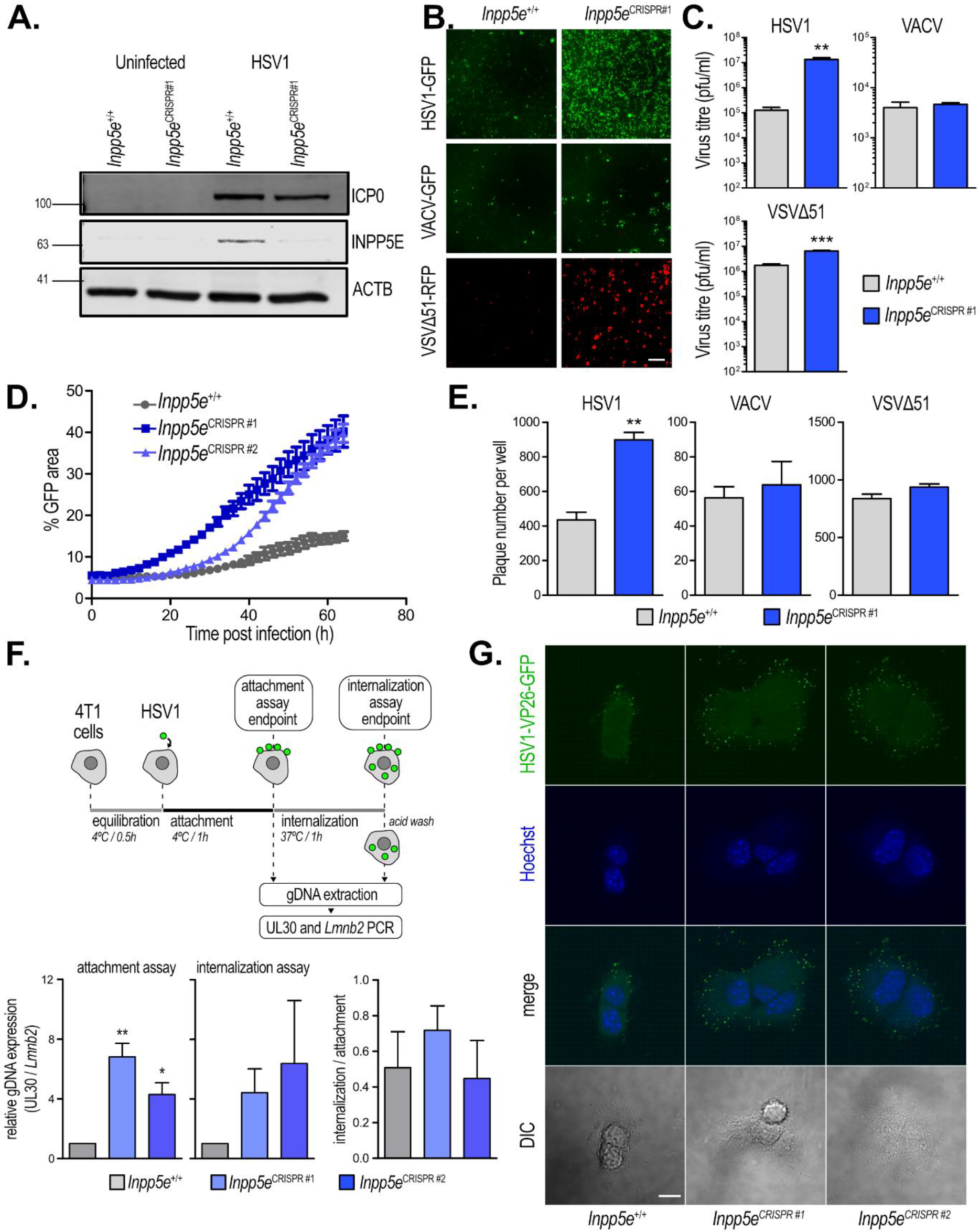
INPP5E functions by impairing HSV1 virion attachment and spread. (**A**) Western blot for HSV1 protein ICP0, INPP5E or β-Actin from lysates 4T1 Inpp5e^+/+^ or Inpp5e^CRISPR#1^ cells. Cells were infected with HSV1-GFP at MOI of 5 for 48 hours. (**B**) Representative fluorescence microscopy images of virus infections at MOI of 0.1 for 24 hours (VSV∆51-RFP) or 48 hours (HSV1-GFP and VACV-GFP) in Inpp5e^CRISPR#1^ vs. Inpp5e^+/+^ 4T1 cells. (**C**) Titration of virus obtained from the supernatant of Inpp5e^CRISPR#1^ vs. Inpp5e^+/+^ 4T1 cells infected with the indicated viruses at 0.01 MOI for 24 hours (VSV∆51-RFP) or 48 hours (HSV1 and VACV). (**D**) Monitoring of HSV1-GFP infection in both Inpp5e^CRISPR^ clones compared to control 4T1 using the Incucyte Live Cell Imaging system. Cells were infected at MOI of 0.1 and images were taken every 2 hours. (**E**) Plaque assay in indicated CRISPR cells. Cells were infected with VSV∆51, HSV1 and VACV at MOI of 0.01, then plaques were enumerated from full-well fluorescence microscopy images. (**F**) Upper panel: Schematic of the cold-binding assays. Inpp5e^CRISPR^ and Inpp5e^+/+^ cells were incubated at 4°C with HSV1 at 10 MOI for 1 hour (attachment assay) or 4°C for 1 hour followed by 37°C for 1 hour (internalization assay). Total DNA was extracted and viral UL30 DNA was quantified by qPCR and normalized to cellular Lmnb2 DNA. Lower panels: Graphs presenting the effect of Inpp5e^CRISPR^ on virus attachment (left), internalization (middle) and the contribution of internalization vs. attachment in each cell type expressed as a ratio (right). (**G**) HSV1-K26-GFP cold-binding assay. As in **F** but using a virus decorated with a GFP-fusion coat protein allowing detection of diffraction-limited puncta by confocal microscopy in fixed cells. Images are single confocal slices representing the mid-cell region. Nuclei are stained with Hoechst. DIC, differential interference contrast. Scale bar: 10 μm

To confirm these results, we quantified the production of infectious virus particles in the media of infected *Inpp5e*^CRISPR^ 4T1 cells. Cells lacking INPP5E exhibited a 2-log increase in HSV1 viral particle production compared to *Inpp5e*^+/+^ 4T1 cells (**Fig. 5C)**. Interestingly, although we observed consistent increases in fluorescence with VSV∆51 by microscopy, there was only a small but significant increase in viral titre (~5-fold increase, p< 0.005) and no significant change was observed in VACV titres, suggesting that INPP5E effect in curtailing infection might be virus-specific. Notably, RNA-mediated reduction in BBS9, the other ciliary gene we identified under translation control, also rendered 4T1 cells more permissive to viral infection (**Fig. S5E, F**), while further decreasing the expression of two translationally downregulated genes, *Usp18* and *Usp44*, compromised slightly HSV1 infectivity (**Fig. S5G, H**).

To determine the stage(s) of the viral life cycle that could be affected by *Inpp5e*, we examined whether *Inpp5e*^CRISPR^ cells could support increased viral spread. We infected cells with HSV1-GFP at low MOI (0.1) and measured GFP intensity and area every 2 hours using a live cell imaging system. By quantitating the area of GFP signal in infected clusters normalized to the total imaged surface area at multiple time points, a more pronounced spread in the two *Inpp5e*^CRISPR^ 4T1 cell lines compared to the control cell line was observed (**Fig. 5D**). As a proxy measurement for viral binding and/or entry, we modified the classical plaque assay by infecting a monolayer of live *Inpp5e*^CRISPR^ or *Inpp5e*^+/+^ 4T1 cells in an agar matrix (to prevent cell-to-cell spread of virion) with HSV1, VSV∆51 or VACV at very low MOI (0.01). *Inpp5e*^CRISPR^ cells generated over 2-fold more plaques when exposed to HSV1 in contrast to no change in the number of VACV or VSV∆51 plaques (**Fig. 5E**). These data suggest that viral binding and/or entry of HSV1 is enhanced in *Inpp5e*^CRISPR^ cells.

To further investigate the role of INPP5E in early HSV1 infection, we separately assessed cell attachment or internalization of virions using two parallel cold-binding assays (**Fig. 5F**, schema in top panel) that quantified virus particles via qPCR of HSV1 genomic DNA (gDNA) copies ^54^. In the attachment assay, 6.8- and 4.3-fold more viral DNA was found associated with the surface-bound fraction in *Inpp5e*^CRISPR#1^ and *Inpp5e*^CRISPR#2^ vs. *Inpp5e*^+/+^ cells, respectively (**Fig. 5F**, lower left panel). A similar trend was measured with the internalization assay, where 4.4- and 6.4- fold more viral DNA was found within *Inpp5e*^CRISPR#1^ and *Inpp5e*^CRISPR#2^ cells respectively, relative to control cells (**Fig. 5F**, lower middle panel). The internalization:attachment ratio did not significantly change in the *Inpp5e*^CRISPR^ cells, indicating that the binding of HSV1 virions to the cell surface is significantly enhanced independently of internalization (**Fig. 5F**, lower right panel). Furthermore, in a parallel cold-binding experiment using HSV1 expressing a GFP-fusion of the capsid protein VP26, which allows visualization of virions as diffraction-limited puncta ^55^, a sizable increase in the number of GFP puncta binding to *Inpp5e*^CRISPR^ cells was observed (**Fig. 5G**).

Together, these data define new translationally regulated antiviral effectors, two of which are the inositol 5-phosphatase INPP5E and the BBSome scaffold protein BBS9. While INPP5E is translationally induced by all three of the viruses we assayed, its expression appears to have a pronounced antiviral effect on HSV1 in 4T1 cells, where it modulates viral attachment and spread.

## DISCUSSION

In response to virus infection, mammalian cells alter mRNA translation, while viruses often attempt to take control of the translation machinery to favour viral mRNA translation ^3,4^. Suppression of global protein synthesis is an innate response to thwart infection. However, synthesis of antiviral proteins is still needed, and antiviral transcripts uniquely upregulated at the translation level, distinct from transcriptionally induced ISGs, have previously been identified by single gene approaches ^56,57^. Our current work reveals a global profile of host cellular mRNAs differentially translated in a murine model of breast cancer cells infected by clinically relevant oncolytic viruses, and it characterizes the regulatory mechanism and function of a new representative translationally induced antiviral gene.

Functional enrichment analysis of translationally regulated mRNAs highlighted genes whose protein products function in primary cilium homeostasis. *INPP5E* is a target of genetic mutation responsible for ciliopathies such as Joubert and MORM syndromes ^58,59^. Mainly localized at the primary cilium, it has also been observed within the nucleus and at centrioles ^60^. Our immunofluorescence data shows virus-induced INPP5E present throughout the cell but with globular foci at perinuclear structures and proximal to the plasma membrane. We observed that cells lacking INPP5E showed increased HSV1 infection and that this can be due to enhanced binding of the virus to the cell surface, while internalization remained unaffected. Importantly, while INPP5E is reported to be localized to the cilium^59^, viral attachment did not appear to be polarized as one would expect if enhanced binding occurred at/or near ciliary structures. Thus, INPP5E antiviral activities might be not related to its cilium localization.

Given that INPP5E is an inositol 5- polyphosphatase, it could conceivably control second messenger phosphatidylinositol (PI) signalling at the plasma membrane as it does within the cilium where it maintains a high concentration of PI-4-phosphate (PI(4)P) relative to PI(4,5)P_2_, an attribute linked to ciliary stability ^61,62^. Indeed, another second messenger, PI(5)P is induced by Newcastle disease virus infection and poly(I:C), and can act as an innate immune effector promoting type I IFN production through the TBK1-IRF3 signalling axis ^63^. Several viruses activate PI3K signalling pathways ^64^; this increases PI(3,4,5)P_3_ and consequentl activates the Akt-mTOR signalling axis, thus modulating infection efficiency. Intriguingly, PI(4,5)P_2_ and PI(3,4,5)P_3_, two likely INPP5E substrates, are well-known mediators of actin remodelling ^65^. This positive effect of INPP5E removal on HSV1 infection may be explained by changes to the actin cytoskeleton (e.g. changes in membrane ruffling that alter viral attachment)^66^. Away from the plasma membrane, centrosomal PIs can also be affected by a lack of INPP5E, leading to spindle microtubule destabilization, possibly through an imbalance in PI(4,5)P2 expression ^60^. Whether these mechanisms have a role to play in mediating the enhanced virus binding and infection that is seen in 4T1 cells is the subject of future investigation.

Another ciliary gene we examined, *BBS9*, encodes a protein that serves as a structural component of the BBSome, a stoichiometric, octameric protein complex (composed of BBS1, 2, 5, 7, 8, 9, 10) charged with receptor cargo destined for the primary cilium ^32^; yet others have shown that the BBSome is important in the transport of extra-ciliary cargo to the plasma membrane ^67^ as well as in retrograde transport ^68^. Mutations in *BBS9* and other *BBS* genes underlie Bardet-Biedel syndrome, a rare, pleiotropic disorder that is thought to arise from cilia dysfunction. Akin to INPP5E, we report here that BBS9 expression is translationally induced during viral infection and limit HSV1, VACV, and VSV∆51 infections. Whether impaired BBSome function is responsible for the proviral effect that we observed with BBS9 knockdown in 4T1 or CT2A cells is unclear, or even if the lack of the BBSome per se is responsible for this effect as BBS complexes lacking BBS9 might display ancillary, proviral functions.

The role of the primary cilium during viral infection of cancer cells requires more investigation. Indeed, understanding how this signalling antenna of the cell might affect viral infection is an intriguing line of study: Are antiviral receptors (e.g. Interferon receptors) trafficked to the plasma membrane or ciliary pocket via the BBSome or other BBS transport complexes? Perhaps the BBSome has been co-opted by viruses to aid in the transport of their protein cargo. Furthermore, whether individuals suffering from ciliopathies present with impaired cell-intrinsic innate immunity is an open question.

We show here that translationally upregulated mRNAs common to all three infections were normally repressed at the level of translation in the uninfected condition. In a search for a mechanistic explanation of this repression-derepression switch, we noted an enrichment of uORFs in their 5’ leaders, a cis-acting sequence element that can confer translational derepression during accumulation of P-eIF2α ^69^. Further investigation using *Inpp5e* 5’ leader reporter assays revealed no uORF-dependent de-repression during infection. Instead, a variant switch during HSV1 infection or upon treatment with the viral mimic poly(I:C) was observed, producing an alternatively transcribed and spliced transcript with a shortened *Inpp5e* 5’ leader. Critically, we found that the spliced 5’ leader intron harbours repressive uORFs and consequently the short *Inpp5e* 5’ leader is a better substrate for translation than its longer, unspliced counterpart. We also found enrichment of binding motif for the splicing factors SRSF1 and SRSF9 in the 5’ leader of translationally regulated genes, which reinforce the suggestive role of alternative splicing in translation regulation of these genes. Thus, our study presents evidence for a regulatory mechanism in which translational output of an antiviral gene is modulated via a transcript variant shift that increases expression of a normally minor variant that harbours a less translationally repressive leader. These findings are in-line with previous studies proposing a role of regulating protein synthesis via transcript variants ^70^ and follows the axiom that alternative transcription and splicing increases transcriptome diversity from a more limited and inflexible genome. Clearly, differences in the coding region between transcript variants can produce functionally different protein isoforms, but differences in the non-coding regions (5’ leader and 3’ UTR) similarly confer important changes to protein function by altering protein abundance in time and space ^15,70^. These regions contain cis-elements that regulate mRNA turnover, location and/or translation. They can be altered via alternative transcription, splicing, start-site selection or alternative polyadenylation and termination. Examples of differential transcript variant expression have been reported during infection with various viruses ^71,72^. More specifically, RNA-Seq has revealed widespread disturbances to host transcription termination caused by HSV1 infection ^39^ and has identified HSV1 ICP27 protein as a major viral factor responsible for modulating the host transcript variant landscape ^38^. Aberrant splicing can also affect innate immunity during host response to virus infection: For instance, a recent report describes that the lack of the RNA binding protein BUD13 induces the retention of the 4^th^ intron in human *IRF7* upon stimulation by poly(I:C) or IFN-α, generating a defective mRNA and impaired antiviral response when challenged with VSV ^73^.

Here, we observed the opposite effect from both a mechanistic and phenotypic perspective: Viral infection or poly(I:C) induced splicing, rather than retention of an intron and a potentiated, rather than abrogated host antiviral response. The question remains how infection signals the switch to an alternative “ribosome-engaged” transcriptome? PKR-mediated splicing has been previously reported ^74^; however, we were unable to link PKR activation to our change in variant expression. Further studies will be required to home-in on the signalling axis that mediates increased expression of the short *Inpp5e* transcript.

In summary, we describe herein a post-transcriptional mechanism for appropriate expression of potent antiviral genes. Our data suggests that innate immunity projects a complicated regulatory landscape in which various host and viral factors are translationally modulated through interactions with pre-mRNA and splicing complexes. In cataloguing the genes that are part of the translational arm of innate immunity, we have uncovered new regulatory nodes that might benefit future study with the goal of improving cancer therapeutics.

## METHODS

### Cell culture and viruses

Mouse breast carcinoma 4T1, NIH3T3, 293T and Vero were obtained from American Type Culture Collection (ATCC). Mouse glioblastoma CT2A was obtained from Dr. David Stojdl (Children’s Hospital of Eastern Ontario Research Institute). PKR-null and respective wildtype MEFs were obtained from Dr. Antonis Koromilas (Lady Davis Institute). Cells were routinely checked for mycoplasma contamination by cytoplasmic DNA staining. NIH3T3, 293T, Vero, CT2A and MEFs were cultured in Dulbecco’s Modified Eagle Medium (DMEM) (HyClone) supplemented with 10% fetal bovine serum (Wisent Bioproducts) and 0.1% penicillin and streptomycin (Life Technologies) at 37°C in 5% CO_2_. 4T1 were cultured in Roswell Park Memorial Institute (RPMI) 1640 (HyClone) supplemented with 10% fetal bovine serum (Wisent Bioproducts) and 0.1% penicillin and streptomycin. HSV1 (HSV1-1716, strain 17 – g34.5 deleted, Sorrento Therapeutics, San Diego, USA), VACV (JX-594 strain Wyeth, Tk-deleted expressing GM-CSF, Jennerex Biotherapeutics / Sillagen, Seoul, Republic of Korea) and Reovirus (Reolysin, Type 3 Dearing, – Oncolytics Biotech, Calgary, Canada) were kindly provided by manufacturers. VSV-∆51-RFP (∆M51 with insertion of RFP marker) was kindly provided by Dr. John Bell (OHRI) ^75^. HSV1-K26-GFP was kindly provided by Dr. Karen Mossman (McMaster University). For propagation of HSV-1716 and HSV-K26-GFP, monolayer of Vero cells was inoculated with viruses at a MOI of 0.5 and cultured for 72 hours at 37°C, 5% CO_2_. Supernatant was clarified by centrifuge at 500 × g for 5 min, then filtered (0.45 μm). Virus particles in the supernatant were separated from cellular debris by ultracentrifugation at 17500 × g for 90 min over a sucrose cushion (36% sucrose, 10 mM HEPES, 150 mM NaCl, 0.1 mM EDTA, pH 7.3). Pellets were resuspended in 1X HNE buffer (10 mM HEPES, 150 mM NaCl, 0.1 mM EDTA, pH 7.3) and stored at − 80°C. VSV-∆51 propagation was adapted from a previously described method ^76^. Briefly, monolayer of Vero cells was inoculated with VSV-∆51 at MOI of 0.01. Approximately 24 hours post inoculation, supernatant was collected and clarified by centrifugation at 500 × g for 5 min. Cleared supernatant are filtered (0.2 μm), then virus particles were pelleted by ultracentrifugation at 28,000 × g for 90 min. Virus particles in the pellet were resuspended in DMEM media and stored at −80°C. JX-594 and Reovirus were used directly from stock provided by manufacturer.

### Lentivirus production and plasmids

The following lentiviral vectors were obtained from Sigma Aldrich: SHC002 (shNTC); TRCN0000080705 (shINPP5E); TRCN0000182387 (shBBS9), TRCN0000030789 (shUSP18), TRCN0000030879 (shUSP44). The lentiviral vectors were co-transfected with the packaging plasmids pLP1, pLP2 and pLP/VSVG (Thermofisher) into HEK293T cells. Lentivirus-containing supernatants were collected at 48 and 72 hours post-transfection and filtered (0.45 μm). The filtrate was applied directly to target cells and integration of the expression cassette was confirmed 72 hours post-transduction by puromycin selection at 2 μg/ml for 4 days. The long mouse *Inpp5e* 5’ leader (nt 1-601) was PCR-amplified from a full-length MGC cDNA clone (Genbank: BC052717; cloneID 6837339 from Dharmacon). This clone is missing the first 31 nucleotides of the annotated long *Inpp5e* mRNA transcript (NM_033134). The 601 nt 5’ leader was cloned into the NotI restriction site of pMC (a kind gift of Dr. Martin Holcik, Carleton University) upstream of the CAT reporter maintaining the same reading frame as the endogenous *Inpp5e* transcript. The short *Inpp5e* leader (found in NM_001290437; nt 1-348) was created from the long-leader CAT construct by a deletion overlap extension cloning strategy. Site-directed mutagenesis was performed to mutate uORF start codons in the long-leader CAT construct. All constructs were verified by Sanger sequencing.

### Polysome profiling

Polysome profiling was conducted as previously described ^77^. Briefly, cycloheximide (Bioshop, CAT #66-81-9) was added directly to the culture media to a final concentration of 100 μg/ml and incubated for 5 min. Cells were then washed 3 times with ice cold PBS containing cycloheximide, then were scraped from the dishes and pelleted at 500 × g for 5 min at 4°C. Cells were lysed with hypotonic buffer (5 mM Tris pH 7.5, 2.5 mM MgCl_2_, 1.5 mM KCl) supplemented with cOmplete Protease Inhibitor Cocktail (Roche) and debris was cleared by centrifugation at 14,000 × g for 5 min, 4°C. Lysates containing ribosome-bound mRNAs were collected, flash frozen, then stored at −80°C. A volume of lysate equal to 10 OD260 units was added on a 10-50% sucrose gradient and centrifuged at 36,000 rpm in a SW41 bucket rotor for 90 min at 4°C. Fractions were collected using a Brandel Fraction Collector System. RNA was extracted from each fraction using Trizol reagent according to the manufacturer’s protocol.

### Metabolic labelling and cell viability

Cells were incubated with complete growth media supplementing with EasyTagTM Express Protein Labeling Mix containing both [^35^S]-L-methionine and [^35^S]-L-cysteine (PerkinElmer) at 10 μCi/ml for 30 min at 37°C, 5% CO_2_. Cells were then lysed in radioimmunoprecipitation assay (RIPA) buffer (150 mM NaCl, 1.0% IGEPAL-CA-630, 0.5% sodium deoxycholate, 0.1% SDS, 50 mM Tris, pH 8.0). Proteins were then separated on SDS-PAGE and transferred to a nitrocellulose membrane, followed by exposure to autoradiography films. Cell viability was measured 48 hours post-infection using the Cell Proliferation Kit I – MTT (Roche) according to manufacturer’s manual as previously described^78^.

### Ribosome profiling

Ribosome profiling was performed as previously described on 2 biological replicates ^79^. Briefly, polysomes in 4T1 lysates were stabilized with cycloheximide and 4T1 lysates were split into two parallel workflows. RNA-Seq on total RNA from one half of the lysate (see below) while RNase I footprinting was performed on the remaining half to capture RPFs. For RNA-Seq, 150 μg of total RNA was taken for RNA-Seq analysis and Poly-(A)+ mRNA was purified using magnetic oligo-dT DynaBeads (Thermofisher) according to the manufacturer’s instructions. Purified RNA was eluted and mixed with an equal volume of 2X alkaline fragmentation solution (2 mM EDTA, 10 mM Na_2_CO_3_, 90 mM NaHCO_3_) and incubated for 20 min at 95°C. Fragmentation reactions were mixed with stop/precipitation solution (300 mM NaOAc pH 5.5 and GlycoBlue), followed by isopropanol precipitation by standard methods. Fragmented mRNA samples were size-selected on a denaturing 10% urea-polyacrylamide gel. The area corresponding to 35-50 nucleotides was excised, eluted and precipitated with isopropanol. Isolated RPF RNA (corresponding to 28-32 nt fragments) and total RNA fragments were used to create cDNA libraries as previously described ^79^. Ribosomal RNA (rRNA) contamination was reduced by subtractive hybridization using biotinylated oligos that were reverse complements of abundant rRNAs. The mRNA and ribosome-footprint libraries were then amplified by PCR (10 cycles) using indexed primers and sequenced on the Illumina HiSeq 2000 platform with read length of 50 nucleotides at the McGill University and Génome Québec Innovation Centre.

### Mapping and analysis of ribosome profiling data

The adapter sequence was removed from reads using FASTX ^80^ and reads that mapped to rDNA sequence by Bowtie 2 ^81^ were discarded. Reads were then mapped to the mouse genome (mm9) using Bowtie 2. Uniquely mapped reads with MAPQ score ≥10 were used for further analysis. For gene expression analysis, reads mapping to the coding region of RefSeq transcripts were used to calculate Reads Per Kilobase per Million total uniquely mapped reads (RPKM). Gene-level RPKMs were obtained by conflating and averaging transcript RPKMs. Genes that showed no expression (0 RPKM) at either the transcription or translation levels in either the mock or infected samples were omitted from further analysis. Translation efficiency was defined by the log_2_ ratio of RPF to total RNA RPKMs. For metagene analysis of read distribution around start and stop codons, reads mapped to RefSeq transcripts were used. For a given region, only genes with at least 128 reads whose 5’ end was within the region were used. The 5’ end position of a read was used for the plotting. Subsets of differentially-expressed genes that are common and unique between the three oncolytic viruses were compiled using Venny v2.1 ^82^.

### Functional analysis of DEGs

Differentially-expressed genes that were commonly up-or downregulated in all three virus infections were tested for enrichment (q<0.10) in Gene Ontology (GO) terms using BiNGO (GO terms downloaded October, 2015) ^83^. GO networks were plotted with the Enrichment Map plugin within Cytoscape 3.0 (www.cytoscape.org). The “R” software package in the RStudio environment or GraphPad Prism was used for all other data manipulation and plotting. For sequence and RNA binding protein analyses, 5’ leaders and 3’ UTRs annotated in RefSeq were retrieved from UCSC Tables.

### Immunocytochemistry

4T1 cells were cultured as indicated on a 10 mm, #1.5 glass cover slip (Electron Microscopy Science). For intracellular staining, cells were washed with ice cold PBS followed by fixation in 4% paraformaldehyde in PHEM buffer (60 mM PIPES, 25 mM HEPES, pH 6.9, 10 mM EGTA, 2 mM MgSO_4_) for 10 min at room temperature. Fixed cells were treated with 50 mM NH_4_Cl in PBS to reduce autofluorescence, followed by permeabilization with 0.1% Triton X-100 in PBS for 10 min. Following 3 washes with PBS, the cover slip was then blocked with 5% BSA in PBS, for 30 min. Permeabilized cells were then incubated with the indicated primary antibody in 1% BSA/PBS overnight. Coverslips were washed 3 times for 5 min each with PBS and incubated with 1:10,000 dilution Goat anti-Rabbit IgG (H+L) Highly Cross-Adsorbed Secondary Antibody Alexa Fluor 568 secondary antibody (#A-11036, Invitrogen) for 1 hour at room temperature in the dark. Coverslips were washed 3 × 5 min with PBS and nuclei were stained with 1:200 of NucBlue^®^ Live Ready Probe-variant of Hoechst dye (Life Technologies) in PBS for 5 min. Cover slips were then mounted onto slides using Prolong^®^ Diamond mounting medium (Life Technologies). Confocal imaging was performed using a FV-1000 confocal microscope (Olympus) with a PlanApo N 100X/1.40, ∞/0.17 oil immersion objective lens (Olympus). Non-confocal fluorescence imaging was performed using an EVOS FL Cell Imaging system (Thermofisher). The following primary antibodies and corresponding dilution are used: 1:100 α-INPP5E (# PA5-37119, Thermofisher), 1:1000 α-BBS9 (#14460-1-AP, Proteintech).

### Western blotting

Cells were washed with PBS and lysed in RIPA buffer supplemented with 10 mM NaF, 10 mM Na_2_VO_3_ and cOmplete Protease Inhibitor Cocktail (Roche). Cells debris was removed by centrifugation at 10,000 × *g*, 10 min, 4°C. Protein concentration of the supernatant was quantified using DC Protein assay (BioRad). Indicated amount of total protein was then loaded onto 10% SDS-polyacrylamide gel and separated by electrophoresis. Separated proteins were transferred to a nitrocellulose or PVDF membrane, then blocked with 5% skim milk in TBS-T buffer (10 mM Tris, 50 mM NaCl, 0.1% Tween-20, pH 7.5). The following primary antibodies and corresponding dilution were used: 1:500 α-INPP5E (# CPA3073, Cohesion Biosciences), 1:10,000 α-β-actin (#A5441, Sigma), 1:5000 α-GAPDH (#8245, Abcam), 1:2000 α- pan-HSV1 (#B011402, Dako), 1:2000 α-HSV1-ICP0 (#11060, Santa Cruz),, 1:10,000 α-β-Actin (#A5441, Sigma Aldrich).

### Quantitative RT-PCR (qRT-PCR) and Droplet Digital RT-PCR (ddRT-PCR)

cDNA was reverse transcribed from total RNA using iScript Advanced cDNA Synthesis Kit (BioRad) according to the manufacturer’s protocol. qRT-PCR was performed on cDNA mixed with iQ SyBR Green mix (BioRad) according to the manufacturer’s protocol, using a Realplex 2 thermocycler (Eppendorf). The PCR conditions were 95°C for 3 min, followed by 40 cycles of 95°C for 10 s, 60°C for 30 s and 72°C for 30 s. Ct threshold was determined by Realplex software (Eppendorf). For calculating mRNA abundance, the ∆∆Ct method relative to *Rps20* expression was used. For calculating short *Inpp5e* variant ratio, the ∆∆Ct method relative to total *Inpp5e* expression was used. For the binding assays, RT-qPCR was performed directly on the extracted DNA using the same mix and PCR conditions as in qPCR assays, then the number of HSV-1 genome relative to the number of host genome was calculated using the ∆∆Ct method comparing relative abundance between HSV-1 *UL30* and mouse *Lmnb2* abundance. For ddRT-PCR, cDNA was mixed with QX200 ddPCR EvaGreen Supermix (Biorad) according to the manufacturer’s protocol and droplets were prepared using QX200 droplet generation oil on a QX200 Droplet Generator (Biorad). Droplets were subjected to PCR using a C1000 thermocycler (BioRad) using the cycling conditions: 95°C for 5 min, followed by 45 cycles of 95°C for 30 s at a ramp rate of 2°C/s and 60°C for 1 min at a ramp rate of 2°C/s. Positive/negative droplets were counted by a QX200 Droplet Reader (BioRad). Primers (listed 5’ to 3’) were used for both qRT-PCR and ddRT-PCR to detect: Total *Inpp5e* (mInpp5e-F: GATCTTTCAGCCTTCTGGCCC, mInpp5e-R: GAGAGCCATGTTTCGGTCTG); short *Inpp5e* (INPP5E-shortUTR-F: CGGAGGGCGCAGGCAT, INPP5E-shortUTR-R: TGAAAACTCGAGTGGCTCCC); *ActB* (mActB-F: GGCTCCTAGCACCATGAAGAT, mActB-R: GGGTGTAAAACGCAGCTCAGTAAC); *Rps20* (RPS20-F: CGCATGCCTACCAAGACTTT, RPS20-R: GGCATCTGCAATGGTGACTTC). For the cold-binding assay, HSV1 gDNA was quantified by qPCR using published primers targeting the UL30 genomic region (HSV-1UL30_F: ACATCATCAACTTCGACTGG, HSV-1UL30_R: CTCAGGTCCTTCTTCTTGTCC) ^84^. Cellular gDNA was quantified using primers targeting *Lmnb2* (m-gDNA-LMNB2-F: ACCAGGTCGTCTGCTATCCT, m-gDNA-LMNB2-R: TCAGTGGTACCTTCAACGCC). For the gel-based assessment of 5’ leader expression, PCR was performed on 4T1 cDNA libraries using INPP5Eutr-F: CAGTCGTTGTTCCAGCTGC and INPP5E-shortUTR-R: TGAAAACTCGAGTGGCTCCC. For normalization of the CAT RNA reporter assay (see below), CAT cDNA was amplified using the above-noted ddPCR conditions with previously published primers ^85^. Melting curves and agarose electrophoresis were performed to control for PCR specificity in all of the above assays.

### CAT translation reporter assays

The CAT assay has been described previously ^12^. Briefly, 4T1, CT2A or MEFs were seeded at 3 or 6×10^5^ cells/well in 6-well plates, then co-transfected with 1 μg each of β-Galactosidase- (pBGal, obtained from Dr. Martin Holcik) and CAT-expressing plasmids using Lipofectamine 2000. 24 hours post-transfection, cells were lysed, and CAT expression was quantified using the CAT ELISA (Roche) as per the manufacturer’s instructions. β-Galactosidase activity was measured using an ortho-Nitrophenyl-β-galactoside (ONPG) colorimetric assay. In the case of RNA transfection experiments, a T7-flanked long *Inpp5e* 5’ leader was amplified from 1 ng of the appropriate CAT reporter plasmid. This amplicon was used as a template for synthesis of capped and poly(A)-tailed RNA using the T7 HiScribe in vitro transcription kit (New England Biolabs) according to the manufacturer’s protocol. 1 μg of RNA was transfected with 2 μl of Lipofectamine 2000, and at 4 hours HSV1 was added into the media at indicated MOIs and incubated for an additional 18 hours. Cells were lysed in 300 μl of CAT lysis buffer, and 30 μl was used to isolate total RNA and prepare cDNA using iScript RT (BioRad). CAT cDNA was amplified by ddPCR using the above-noted primers and conditions. CAT expression was determined as above and normalized to the CAT RNA levels.

### CRISPR/Cas9-mediated gene knockout

CRISPR/Cas9 knockout of INPP5E in 4T1 was performed as previously described ^52^. Briefly, small guide RNAs (sgRNA) targeting the *Inpp5e* first exon (mInpp5e-sgRNA1-F: 5’-CACCGAGCTTGCCTGCGTCACACTG-3’; mInpp5e-sgRNA1-R: 5’-AAACCAGTGTGACGCAGGCAAGCTC-3’) or non-targeting sgRNA (mNT-sgRNA-F: 5’-CACCGCGAGGTATTCGGCTCCGCG-3’; mNT-sgRNA-R: 5’-AAACCGCGGAGCCGAATACCTCGC-3’) were cloned into the lenti-sgRNA(MS2)-zeomycin backbone (Addgene #61427) using BsmBI. To produce separate Cas9- and sgRNA-expressing lentiviruses, doxycycline-inducible and puromycin expressing pCW-Cas9 (Addgene #50661) or the constitutively expressing sgRNA vector were co-transfected with pLP1, pLP2 and pLP/VSVG (Thermofisher) into HEK293T cells using Lipofectamine 2000 (Invitrogen) according to the manufacturer’s protocol. Cells were then transduced with the lentiviral supernatants and double-transductants were selected using puromycin and zeomycin. Cas9 expression was then induced using 1 μg/ml doxycycline for 24 hours. Single-cell clones were obtained by limiting dilution, screened using T7 Endonuclease I (New England Biolabs), and confirmed by Sanger sequencing. CRISPR/Cas9 modification efficiency was quantitated using TIDE, which deduces the frequency of individual insertion-deletion (indel) from Sanger sequencing of a mixed population.

### Live cell monitoring of virus spread

Live cells monitoring of virus infection was performed using the IncuCyte Live-Cell Analysis system (Sartorius). Cells were seeded in a 24-well plated at 80% confluency, then infected with viruses at the indicated MOI. Multiple phase contrast and fluorescence images were taken per well every 2 hours. Images were then analyzed using the IncuCyte Zoom software (Sartorius) for GFP cluster integrated intensity (Green calibrated unit × μm^2^) as a measurement for virus infection. GFP cluster integrated intensity was calculated using the following customized process definition in Incucyte ZOOM software: background subtraction using Top-Hat method (disk shape structuring element with radius of 10 μm, threshold of 1.0 green calibrated unit), edge split: Off, Hole Fill: No, Adjust Size: No, Filters: No.

### Plaque assays

For titration, Vero cells were cultured to a monolayer. Cells were then incubated with a serial dilution in DMEM of virus-containing supernatant for 1 hours, 37°C, 5% CO_2_ with shaking every 10 min. Cells were then washed 3 times with DMEM, then a layer of DMEM + 10% FBS +1% agar was added on top and allowed to solidify. Cells were then cultured at 37°C, 5% CO_2_ for 48 hours (VSV-∆51) or 72 hours (HSV-1, VACV). Full-well fluorescence images were taken, and fluorescent plaques were counted and used for back calculating original viral titer. For modified plaque assay to compare viral entry and spread (**Figure 5F**), monolayers of 4T1 WT or 4T1 Inpp5e^CRISPR^ were cultured, then incubated with 0.01 pfu of viruses per cells (MOI of 0.01) diluted in RPMI media. Cells were then washed 3 times with RPMI media, then a layer of RPMI + 10% FBS +1% agar was added on top and allowed to solidify. Cell culture, and plaque detection and counting were carried out as described above for standard plaque assays.

### Binding assays

The cold binding assay was adapted for HSV1 from a previously described method ^86^. Briefly, cells were pre-incubated at 4°C for 30 min, then incubated with HSV-1716 at the indicated MOI at 4°C for 1 hour. For internalization, cells were incubated for 1 hour at 37°C, then washed with PBS pH 3.0 to remove surface bound viruses. gDNA was then harvested using the QIAamp DNA Mini Kit (Qiagen) according to manufacturer’s protocol. PCR was performed using primers described above on the DNA.

### Statistical analyses

All experiments were performed with at least 3 biological replicates unless otherwise specified. Statistical significance was *a priori* set to 0.05. Two-tailed Student’s t-tests or one-way ANOVAs with Dunett’s post-hoc tests were performed where applicable unless otherwise indicated in the figure legend. Error bars indicate standard error of the mean (sem). *p<0.05, **p<0.01, ***p<0.001, ****p<0.0001, ns, non-significant.

## DECLARATIONS

### Ethics approval and consent to participate

Not applicable

### Consent for publication

Not applicable

### Availability of data and material

The data is available is available from the corresponding author upon reasonable request.

### Competing interests

The authors declare that they have no competing interests.

### Funding

HDH. is a recipient of a University of Ottawa Graduate Scholarship, Ontario Graduate Scholarship, and University of Ottawa Faculty of Medicine Destination 2020 Scholarship. This work was supported in part by the Canadian Breast Cancer Foundation/Canadian Cancer Society Research Institute, the Cancer Research Society/CIHR Institute of Cancer Research (grant #22124), the Natural Sciences and Engineering Research Council of Canada (RGPIW-2016-05228), and the Terry Fox Research Institute (New Frontier Program Project Grant) to T.A. and T.E.G.

### Authors’ contributions

TA, HDH, TEG, SMJ conceived and designed experiments. HDH and TEG performed the majority of experiments. HDH, TEG and TA wrote the manuscript. JJ produced viruses and performed viral titre assays. NV performed heterologous reporter assays. VG performed Western Blot, TA, CGG and SMJ performed the wet component of ribosome profiling. WL performed the ribosome profiling mapping and differential expression analysis. MJ and TA reviewed and edited the final manuscript. All authors read and approved the final manuscript.

## ACKNOWLEDGEMENTS

We thank Drs. John Bell, David Stojdl, Martin Holcik, Antonis Koromilas and Karen Mossmann for experiment materials.

---

**Supplementary Figure 1.**
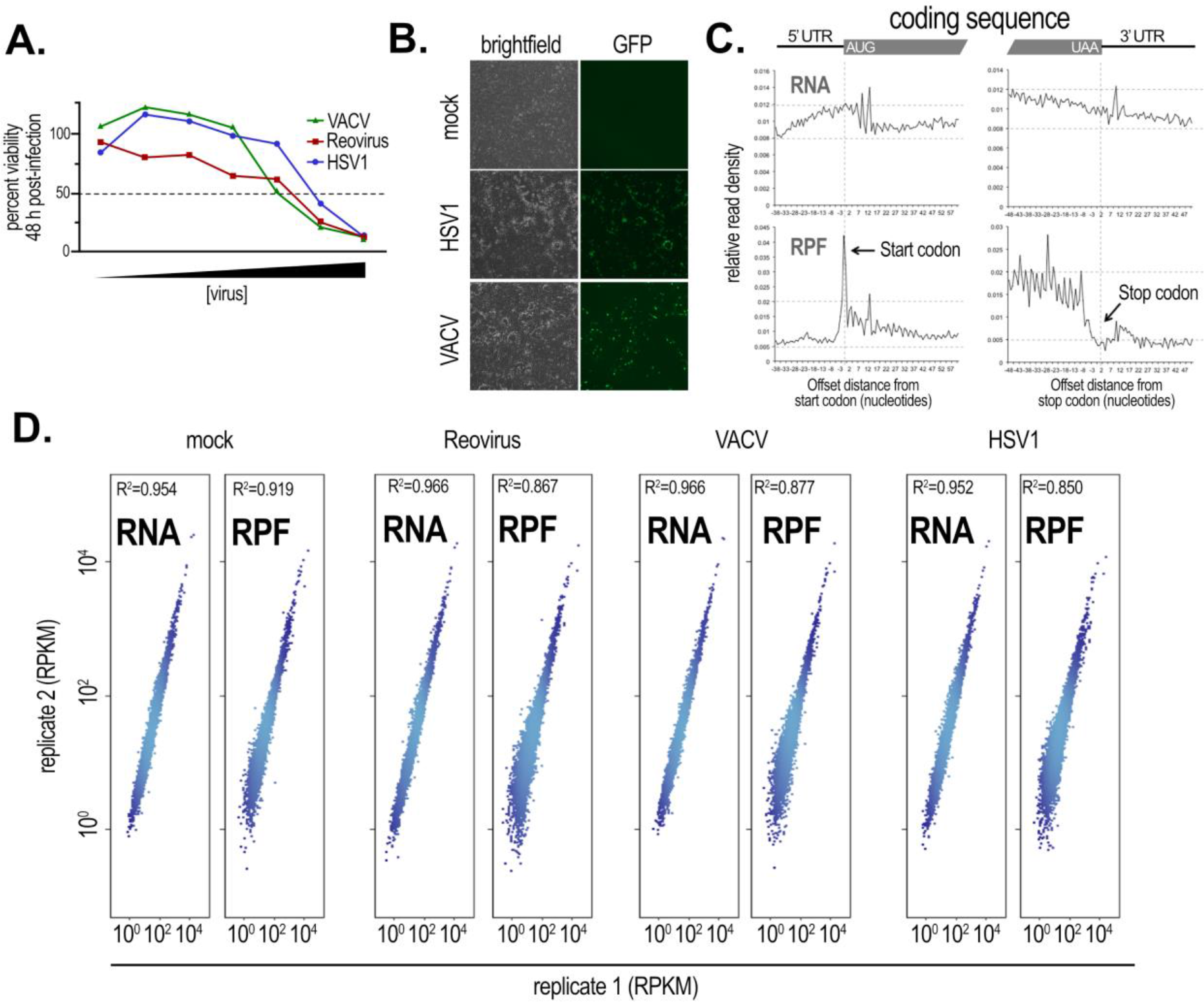
Quality control of the ribosome profiling experiment. **(A)**The effective dose of Reovirus, HSV1 and VACV that reduces the viability of murine 4T1 breast carcinoma cells by 50% after 48 hours of infection (ED_50_) was used to normalize the different infection kinetics of the three viruses prior to ribosome profiling. (**B**) Fluorescence microscopy of 4T1 cells mock-infected or infected with an ED_50_ of HSV1 or VACV (note that Reovirus does not express a fluorescent reporter). (**C**) Metagene analysis of total RNA read (top panels) and RPF read densities (bottom panels) across a randomized selection of mRNAs demonstrates the ability of the ribosome profiling technique to detect RPFs enriched in the coding sequence of mRNAs. (**D**) Scatter plots and correlation analyses of RNA- and RPF-Seq gene expression between biological replicates show a high degree of reproducibility. Linear regression was performed on biological replicate 2 vs 1 and the closeness of fit (R^2^) to the line of regression is reported, where 1.0 is a perfect fit.

**Supplementary Figure 2:**
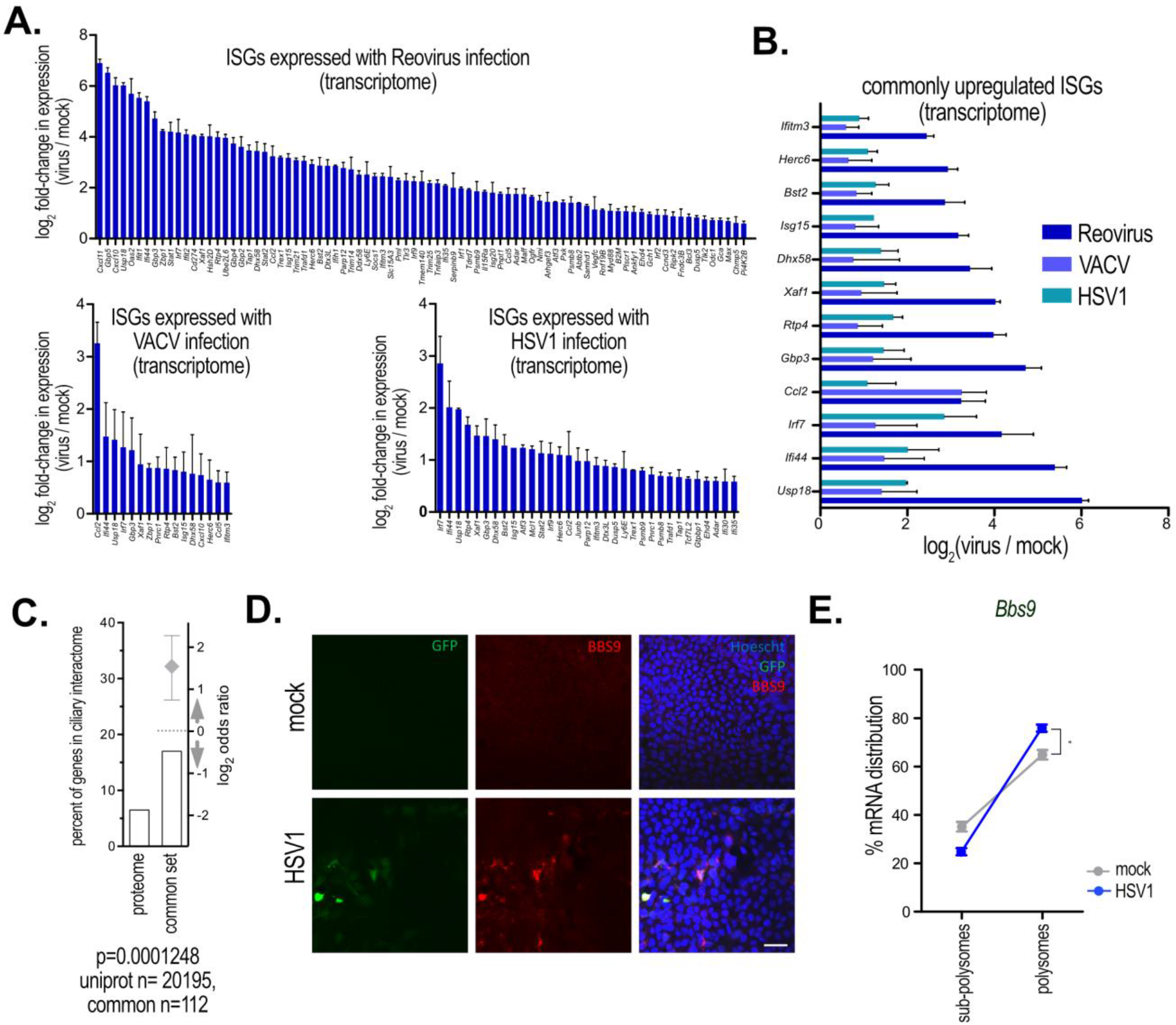
Transcription and translation responses capture an antiviral state. (**A**) Previously validated ISGs ^1^ (from a list of 389) found to be upregulated (>2-fold change relative to mock-infected cells) in the transcriptome of 4T1 cells infected with Reovirus, VACV, or HSV1. (**B**) ISGs from **A**which were found to be transcriptionally regulated in all three infections. (**C**) Frequency of mouse ciliary proteins (based on the human orthologues forming the ciliary interactome ^31^ encoded by the genes in the common up *and* down translationally regulated lists (19/112) vs. the human proteome (1319/20195). (**D**) Immunofluorescence microscopy images of fixed, mock- or HSV1-infected 4T1 cells probed with anti-BBS9 (middle panels). The virus expresses GFP (left panels), and nuclei are stained with Hoechst. **(E)**TE of *Bbs9* increases following infection with HSV1 in 4T1 cells (left panel) as assessed by analyzing the mRNA distribution by polysome profiling (right). Two biological replicates are plotted.

**Supplementary Figure 3:**
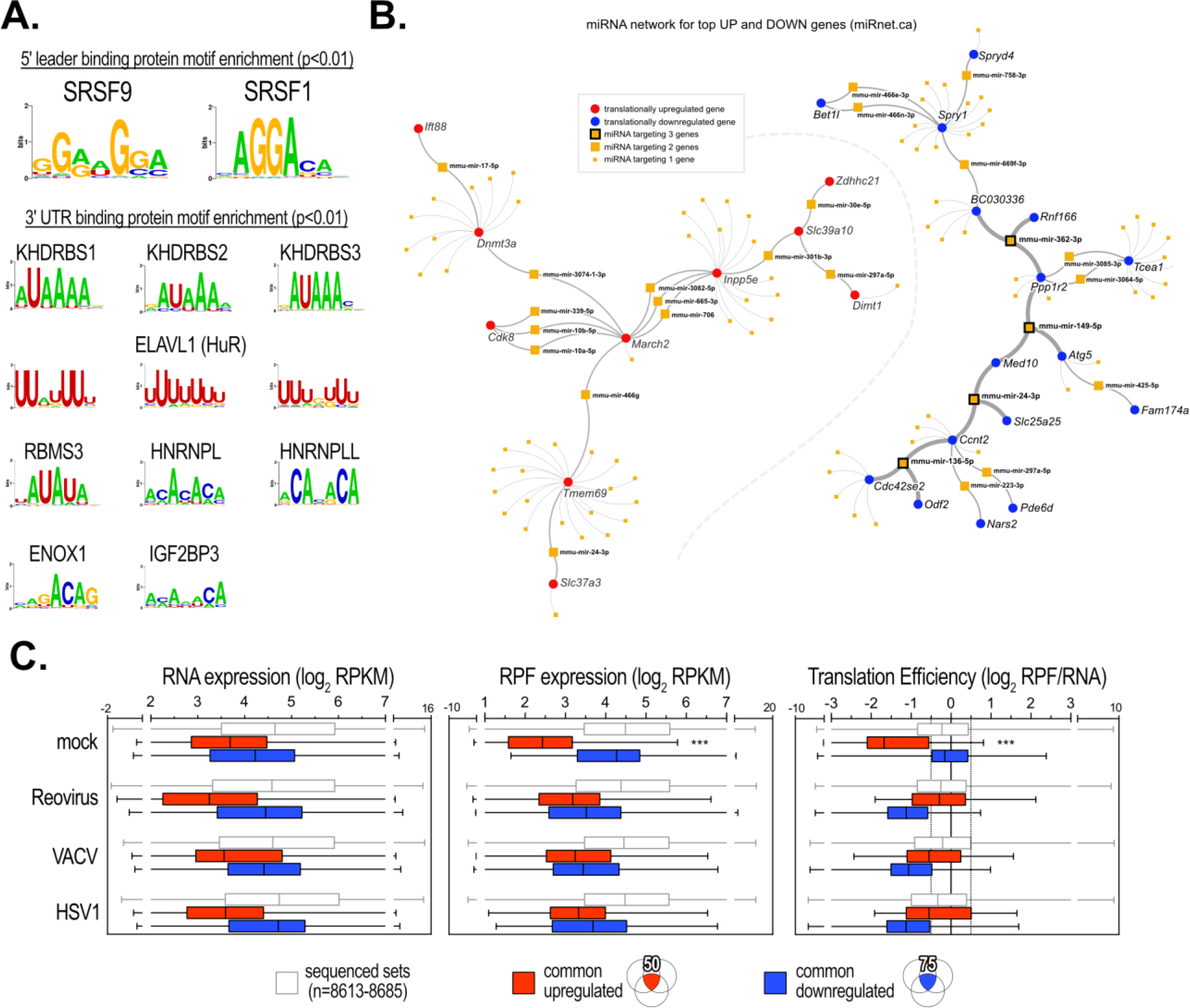
Characterizing cis-elements in the common translationally regulated transcripts. Sequence logos of RNA binding protein motifs enriched in 5’ leaders and 3’ UTRs of transcripts comprising the commonly translationally up-*and* down-regulated list. The CISBP-RNA database^36^ was queried using the Analysis of Motif Enrichment (AME), part of the MEME suite^35^ **(B)**miRNA target analysis was performed using miRNet^40^ with the common translationally up *or* down gene lists and plotted as a network using the built-in network visualizer. (**C**) RNA, RPF expression and TE in each condition for all genes, common translationally upregulated, or common translationally downregulated genes. Translationally upregulated genes represent a rare subset of mRNAs that are translationally repressed under basal conditions in 4T1 cells (c.f. mock expression in RNA and RPF plots).

**Supplementary Figure 4:**
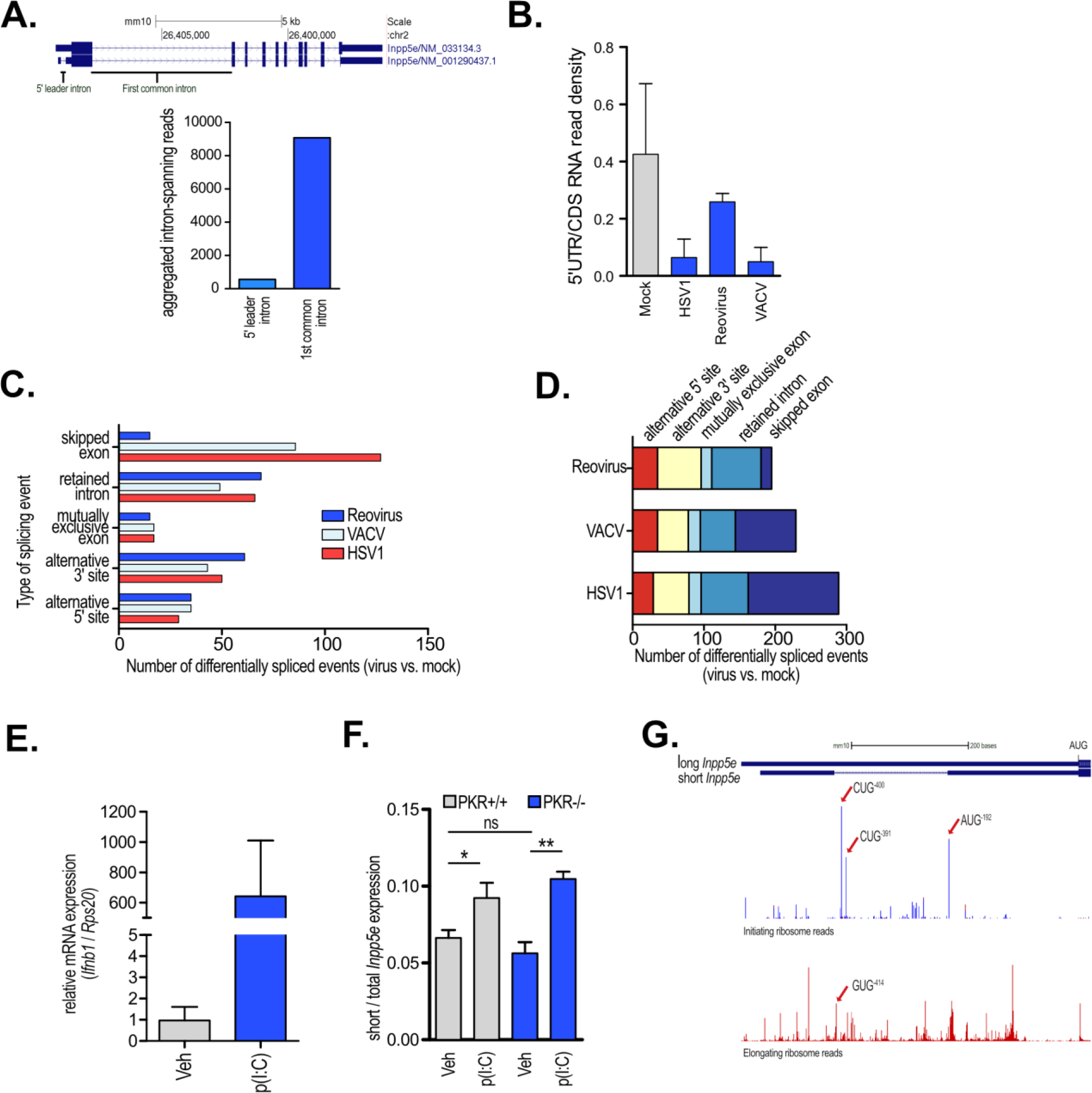
Evidence for*Inpp5e* variant switching with infection or poly(I:C) treatment. (**A**) Model of Refseq-annotated transcript variants of *Inpp5e* (upper illustration) and graph (lower panel) representing the number of intron-spanning reads associated with a pool of samples from the sequencing read archive (NCBI Mus musculus Annotation Release 106, “RNA-Seq intron-spanning reads, aggregate”) from the intron residing in the 5’ leader and the first intron common to both transcript variants. (**B**) Coverage ratio of RNA-Seq reads mapping to the 5’ leader vs. CDS in mock- or virus-infected 4T1 cells. (**C**) MISO analysis detects differentially spliced events in the transcriptome of 4T1 infected with Reovirus, VACV or HSV1, grouped by event types. (**D**) Same analysis as in **C**but with events classified by viral infection. (**E**) qRT-PCR of Ifnb1 normalized to Rps20 CT2A cells treated with poly(I:C) at the indicated times. Data are from 4 independent experiments. (**F**) Poly(I:C)-mediated induction of the short *Inpp5e* variant is PKR- independent. 8 μg/ml of poly(I:C) was transfected into PKR^+/+^ or PKR^−/−^ MEFs and short vs. total *Inpp5e* RNA levels were determined by qRT-PCR 18 hours later. (**G**) Genome Wide Information on Protein Synthesis (GWIPS) view of the 5’ leader region of *Inpp5e* (top) showing initiating ribosome read density (middle; densities are aggregated from all ribosome profiling experiments that assay ribosomes at initiation codons in the database) and elongating ribosome reads (bottom). Red arrows mark initiating or elongating ribosome peaks corresponding to the indicated putative uORF start codon.

**Supplementary Figure 5:**
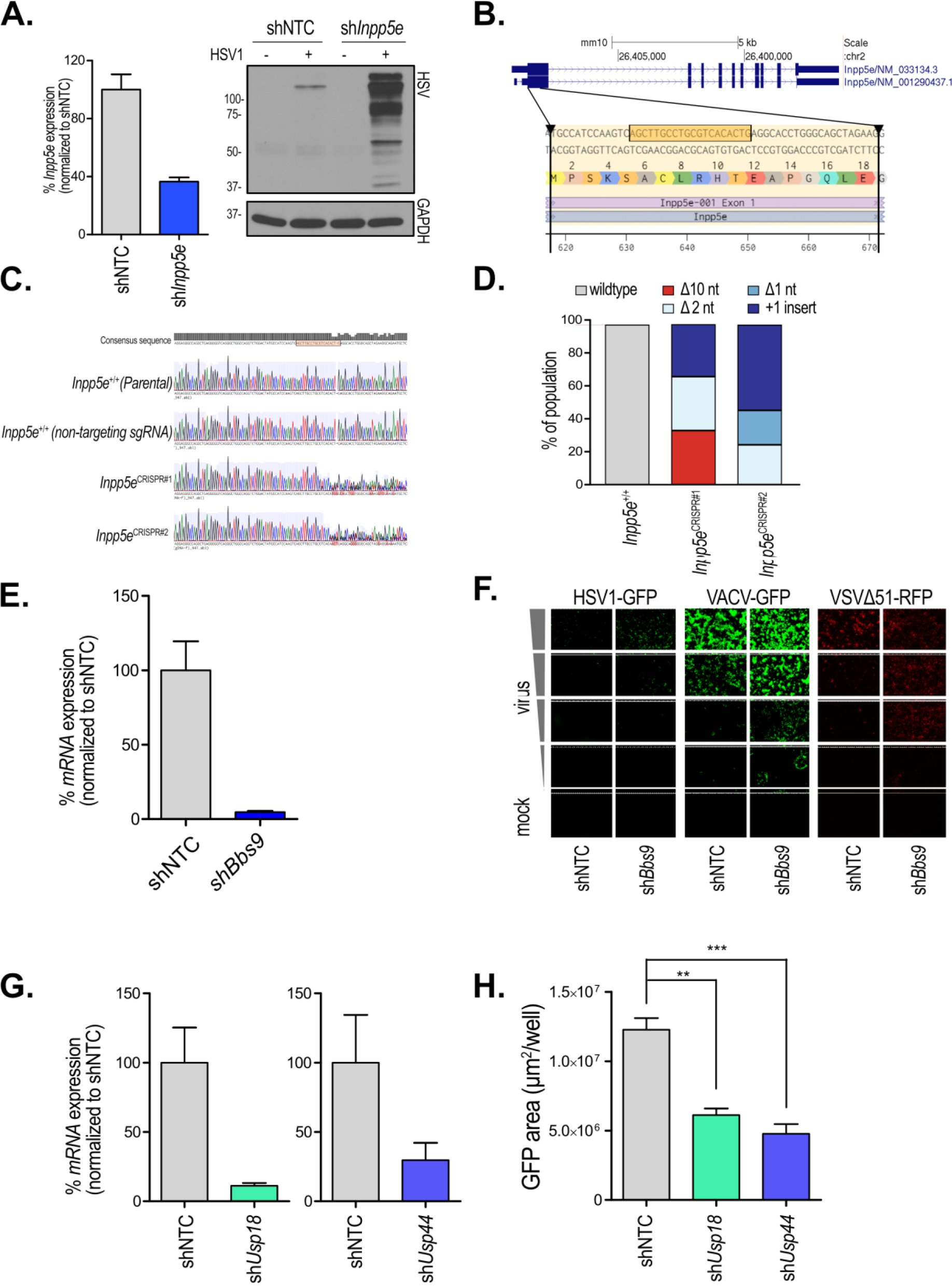
Characterizing the role of candidate proteins in modifying viral infection. (**A**) Knockdown efficiency using shRNA targeting *Inpp5e* vs. non-targeting control (shNTC) in CT2A cells assessed by qRT-PCR. Western blot of lysates from shNTC vs. *Inpp5e* knockdown CT2A cells infected with HSV1 at a MOI of 0.1 for 48 hours for HSV1 proteins or GAPDH. **(B)**Targeting region and sequence of the synthetic single guide RNA (sgRNA) used to disrupt both *Inpp5e* variants. (**C**) Sanger sequencing chromatograms of the wildtype and two *Inpp5e*^CRISPR^ clones. The chromatogram begins at the first nucleotide of the sgRNA sequence. PHRED quality scores are indicated at the top of each chromatogram. (**D**) Indel frequency of the two *Inpp5e*^CRISPR^ clones evaluated using the sequence decomposition program TIDE. Types of indel are categorized and plotted as a percentage of the entire population. (**E**) Knockdown efficiency of *Bbs9* as evaluated by qRT-PCR. (**F**) Representative immunofluorescent images of 4T1 cells stably expressing indicated shRNA (a scrambled control, shNTC; and two different shRNAs targeting *Bbs9*, sh*Bbs9-1* and *-2*) and infected with indicated viruses at increasing MOI. (**G**) Knockdown efficiency of *Usp18* (left) and *Usp44* (right) in 4T1 cells using two different shRNA sequences for each target and assayed by qRT-PCR. (**H**) Infectivity of HSV1-GFP in 4T1 cells stably expressing shRNA targeting *Usp18* and *Usp44*, as measured by GFP-expressing area

